# Aberrant oxidative metabolism selects for *TET2*-deficient hematopoietic stem and progenitor cells

**DOI:** 10.64898/2026.02.28.707294

**Authors:** Katia E. Niño, Vera Adema, Alyx E. Gray, Courtney M. Cowan, Wolfgang E. Schleicher, Mohsen Hosseini, Sierra N. Bennett, Sweta B. Patel, Steven Moreira, Etienne P. Danis, Feiyang Ma, Hsin-Ying Lin, Tracy N. Young, Colin C. Anderson, Devyani Sharma, Angelica Varesi, Marie-Dominique Filippi, Keisuke Ito, Meelad M. Dawlaty, Gang Huang, Julie A. Reisz, Stephanie Z. Xie, Steven M. Chan, Lin Tan, Guillermo Garcia-Manero, Kelly S. Chien, Irene Gañan Gomez, Angelo D’Alessandro, Simona Colla, Eric M. Pietras

**Affiliations:** Division of Hematology, Department of Medicine, University of Colorado Anschutz Medical Campus. Aurora, Colorado, USA 80045; Department of Leukemia, The University of Texas MD Anderson Cancer Center. Houston, Texas, USA 77030; Department of Biomedical Informatics, University of Colorado Anschutz Medical Campus. Aurora, Colorado, USA 80045; University of Colorado Cancer Center, University of Colorado Anschutz Medical Campus. Aurora, Colorado, USA 80045; Princess Margaret Cancer Centre, University Health Network, Toronto, Ontario, Canada; Department of Medical Biophysics, University of Toronto, Ontario, Canada; Department of Biochemistry and Molecular Genetics, University of Colorado Anschutz Medical Campus. Aurora, Colorado, USA 80045; Division of Experimental Hematology and Cancer Biology, Cincinnati Childrens’ Hospital Medical Center. Cincinnati, OH USA 45267; Departments of Cell Biology, Medicine and Oncology, Ruth L. and Davis S. Gottesman Institute for Stem Cell and Regenerative Medicine Research, Albert Einstein College of Medicine. Bronx, New York, USA 10461; Departments of Genetics and Developmental and Molecular Biology, Ruth L. and Davis S. Gottesman Institute for Stem Cell and Regenerative Medicine Research, Albert Einstein College of Medicine. Bronx, New York, USA 10461; Department of Cell Systems and Anatomy, University of Texas at San Antonio. San Antonio, Texas USA 78229; Department of Bioinformatics and Computational Biology, The University of Texas MD Anderson Cancer Center. Houston, Texas USA 77030; Department of Microbiology and Immunology, University of Colorado Anschutz Medical Campus. Aurora, Colorado, USA 80045

## Abstract

The mechanism(s) driving selective expansion of mutant hematopoietic stem and progenitor cells (HSPC) in clonal hematopoiesis (CH) are incompletely understood. Here, we address the role of metabolism in selection for HSPC with loss of function mutations in *TET2*. Loss of *Tet2* in murine HSPC triggers overexpression of glycolysis and oxidative phosphorylation genes and increased oxidative metabolism via an enlarged mitochondrial network. However, *Tet2*-deficient HSPC maintain a normal redox state. Strikingly, compound loss of the rate-limiting pentose phosphate pathway (PPP) enzyme glucose-6-phosphate dehydrogenase (G6PD) triggers increased reactive oxygen species and impairs the fitness of *Tet2*-deficient HSPC. We find that aberrant oxidative metabolism is also a feature of HSPC in human CH and clonal cytopenia of unknown significance (CCUS). Overall, our data point to aberrant metabolism as a critical and conserved driver of selection in *TET2*-deficient CH and identify the PPP as a crucial compensatory pathway needed to maintain their selective advantage.

**Statement of Significance:** This study identifies oxidative metabolism as a critical driver of selection for *TET2*-deficient HSPC in clonal hematopoiesis (CH). It also demonstrates that cellular redox state is a vulnerability that impairs their fitness. These insights establish targetable metabolic pathway(s) that could be exploited in the setting of *TET2* mutant CH.

## Introduction

Clonal hematopoiesis (CH) is the culmination of somatic evolution in the bone marrow (BM), resulting in over-representation of hematopoietic stem and progenitor cell (HSPC) clones that harbor leukemia-associated mutations (1-3). These clonal fractions contribute to risk for a spectrum of malignant and non-malignant diseases including blood and solid tissue cancers, cardiovascular disease, and other conditions that overall contributes to increased risk of all-cause mortality (4-12). Given the prevalence of CH in aged populations over 55 years of age can reach well above 20%, with increased risk of morbidity and mortality for those with mutant variant allele frequencies (VAF) of >10%, the scientific and translational importance of characterizing the mechanism(s) driving selection for CH clones is significant given an aging worldwide population (10,13-15). However, these mechanisms are incompletely understood, constituting a roadblock to improved management of CH and its comorbidities.

Approximately 10-30% of CH cases are associated with mutations in *TET2*, which encodes a dioxygenase that catalyzes the hydroxylation of 5-methylcytosine (5mC), the initial reaction required for cytosine demethylation(12,16). This results in patterns of DNA hypermethylation and enhanced self-renewal of *TET2* mutant HSPC (17-19). Furthermore, *TET2* mutant CH leads to production of myeloid cells with enhanced inflammatory potential and is closely associated with a pantheon of inflammatory pathologies including cardiovascular disease (CVD), which remains a leading cause of death with 1 in 5 deaths attributable to CVD in the United States (20-24). Previous studies have identified cell-intrinsic alterations in the PU.1 regulatory network and other transcriptional regulators controlling HSPC self-renewal and differentiation gene programs as hallmarks of *TET2* deficiency that trigger enhanced self-renewal capacity and promote leukemogenesis (25-27). How these molecular changes translate into cellular mechanisms that directly control fate choice remains largely unexplored. We previously showed that loss of PU.1 network activity in a mouse model carrying a hypomorphic allele of the PU.1 - 14kb upstream regulatory element triggers increased expression of genes related to cellular metabolism in PU.1-deficient HSPCs, serving as a potential mechanism facilitating their increased proliferative rate under conditions of inflammatory stress (28,29). In addition, these cells overexpressed compensatory antioxidant gene programs that suppress reactive oxygen species (ROS), thereby limiting ROS-induced cellular differentiation and promoting stemness (28). Based on these findings we hypothesized that loss of *TET2* could license the activation of metabolic gene programs that drive expansion of *TET2*-deficient HSPC while maintaining their enhanced self-renewal capacity.

Here, we address this hypothesis by analyzing the metabolic and gene expression features of HSPC from mice heterozygous for a *Tet2* knockout allele(30), thereby mimicking the genetic features of human CH. Strikingly, we find that *Tet2*-deficient HSPC exhibit aberrant oxidative metabolism and overexpress gene programs associated with glycolysis and mitochondrial oxidative phosphorylation (oxphos). Importantly, we show that increased oxidative metabolism triggers dependence of *Tet2*-deficient HSPC on the pentose phosphate pathway (PPP) for redox homeostasis. Lastly, we show that aberrant oxidative metabolism is likewise a feature of HSPC from patients with clonal cytopenia of unknown significance (CCUS). Taken together, our data show that aberrant oxidative metabolism functions as both a mechanism to facilitate selective expansion of *TET2*-deficient HSPC, as well as a potential vulnerability that could be exploited to limit their enhanced functional potential.

## Results

### *Tet2*-deficient HSPC selectively expand *in vivo* in a non-conditioned mouse model of CH

To model clonal hematopoiesis *in vivo*, we employed a mouse model of CH in which unfractionated wild-type (*Tet2^+/+^*) or heterozygous (*Tet2^+/-^*) bone marrow cells from a well-characterized germline *Tet2* knockout model (30) are adoptively transferred into non-conditioned CD45.1^+^ recipient mice (**Fig. 1A**). This approach allows us to evaluate the dynamics underlying selection for CH HSPC in native BM conditions and in a genetic context that resembles human CH, in which a single allele of TET2 is typically mutated. Using this method, we previously observed stable long-term donor chimerism (22) and here likewise identified a gradual selection for *Tet2^+/-^* donor hematopoiesis by 10 weeks post transfer, characterized by expansion of *Tet2^+/-^* myeloid and lymphoid compartments in the peripheral blood (PB) and moderate neutrophilia (**Fig. 1B-E**, **Fig. S1A-D**). Likewise, by 10 weeks post transfer we observed a significant increase in *Tet2^+/-^* donor chimerism in bone marrow myeloid and HSPC compartments, including granulocyte-macrophage progenitors (GMP), and all three subsets of lineage-primed multipotent progenitor (MPP) (**Fig. 1H-I**, **Fig. S1E-F**) (31-33). On the other hand, we did not yet observe expansion of phenotypic HSC at this timepoint, likely owing to their slow rate of expansion under native conditions (**Fig. 1J**) (22). Taken together, these data show that heterozygous*Tet2*-deficient HSPC selectively expand under native BM conditions.

**Figure 1.**
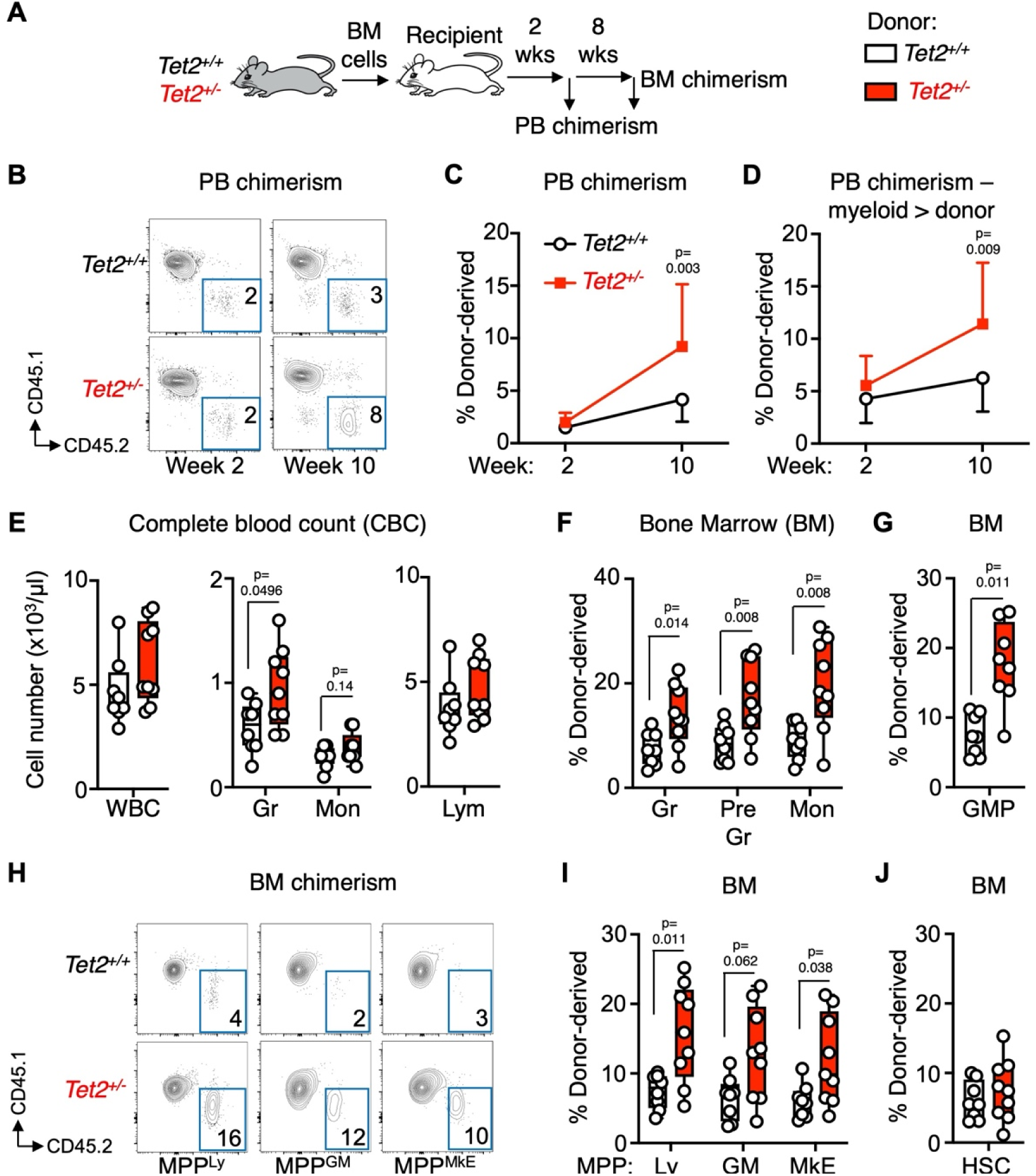
*Tet2*-deficient HSPC selectively expand in a non-conditioned mouse model of CH. Analysis of donor peripheral blood (PB) and bone marrow (BM) chimerism at 2 and 10 weeks post BM adoptive transfer (n = 8 *Tet2^+/+^,* 9 *Tet2^+/-^*). **A**) Study design; **B**) representative flow cytometry plots showing donor and recipient PB chimerism at 10 weeks post adoptive transfer; line graphs showing **C**) donor chimerism and **D**) donor chimerism in the CD11b^+^ myeloid compartment at 2 and 10 weeks post adoptive transfer. **E**) Complete blood count (CBC) analysis of total white blood cells (WBC), neutrophils (Neu), monocytes (Mon) and lymphocytes (Lym) at 10 weeks post adoptive transfer. **F**) Donor chimerism in BM granulocyte (Gr), pre-granulocyte (Pre Gr) and monocyte (Mon) compartments; and **G**) BM granulocyte macrophage progenitor (GMP) compartment. **H**) Representative FACS plots showing donor and recipient multipotent progenitor (MPP) subset chimerism; **I**) donor chimerism in lymphoid (Ly), granulocyte/macrophage (GM) and megakaryocyte/erythroid (MkE)-enriched MPP fractions; and **J**) hematopoietic stem cell (HSC). Error bars in C and D show mean ± SEM. Boxplots show means and individual datapoints and whiskers show minimum/maximum values. Significance was determined via Mann-Whitney *u*-test and exact *p*-values are shown. Data are compiled from two independent experiments.

### *Tet2*-deficient HSPC exhibit aberrant metabolic gene expression programs

To identify the mechanistic basis for the selective expansion of heterozygous *Tet2-*deficient HSPC, we performed single-cell RNA sequencing (scRNA-seq) analysis of *Tet2^+/+^* and *Tet*^+/-^ BM cells. To ensure sufficient representation of rare HSPC populations in our data, we harvested BM from *Tet2^+/+^* and *Tet*^+/-^ mice, sorted Lineage^-^/c-Kit^+^/Sca-1^+^ (LSK) and Lineage^-^/c-Kit^+^ (LK) cell fractions, and recombined these with whole bone marrow (WBM) at a 3:1:1 (LSK:LK:WBM) ratio (**Fig. 2A**). We identified 29 total unique cell clusters (**Fig. S2A, Table S1**). We did not identify any novel or missing clusters between genotypes and observed relatively similar proportions of cell types (**Fig. 2C**). Differential expression analysis of HSPC populations revealed numerous significantly differentially regulated genes in each population (**Fig. 2D, Table S2**). Fast Gene Set Enrichment Analysis (fGSEA) of key HSPC populations revealed upregulation of interferon and mTOR gene signatures in *Tet2^+/-^* HSPC, as well as enrichment of glycolytic, hypoxia and oxidative phosphorylation (oxphos) genes, particularly in MPP and GMP (**Fig. 2E-H**, **Fig. S2C-D**, **Table S3**). To validate our findings in an independent gene expression dataset, we reanalyzed publicly available bulk RNA-seq analysis (GSE132090) (27) of LSK cells isolated from *Tet2^-/-^* mice generated using a different genetic targeting strategy (34), as well as from *Tet2^mut^*mice bearing two homozygous amino acid substitutions (H1367Y and D1369A) that deactivate the enzymatic activity of *Tet2*. Differential expression and GSEA enrichment analysis showed similar patterns of enrichment for Oxphos and glycolysis genes (**Fig. S2E-F**, **Table S3**) in both *Tet2*-deficient models, with substantial overlap in differentially expressed genes comprising both signatures (**Fig. S2F**). Collectively, these data uncover broad overexpression of oxidative metabolism gene programs in heterozygous *Tet2*-deficient HSPC.

**Figure 2.**
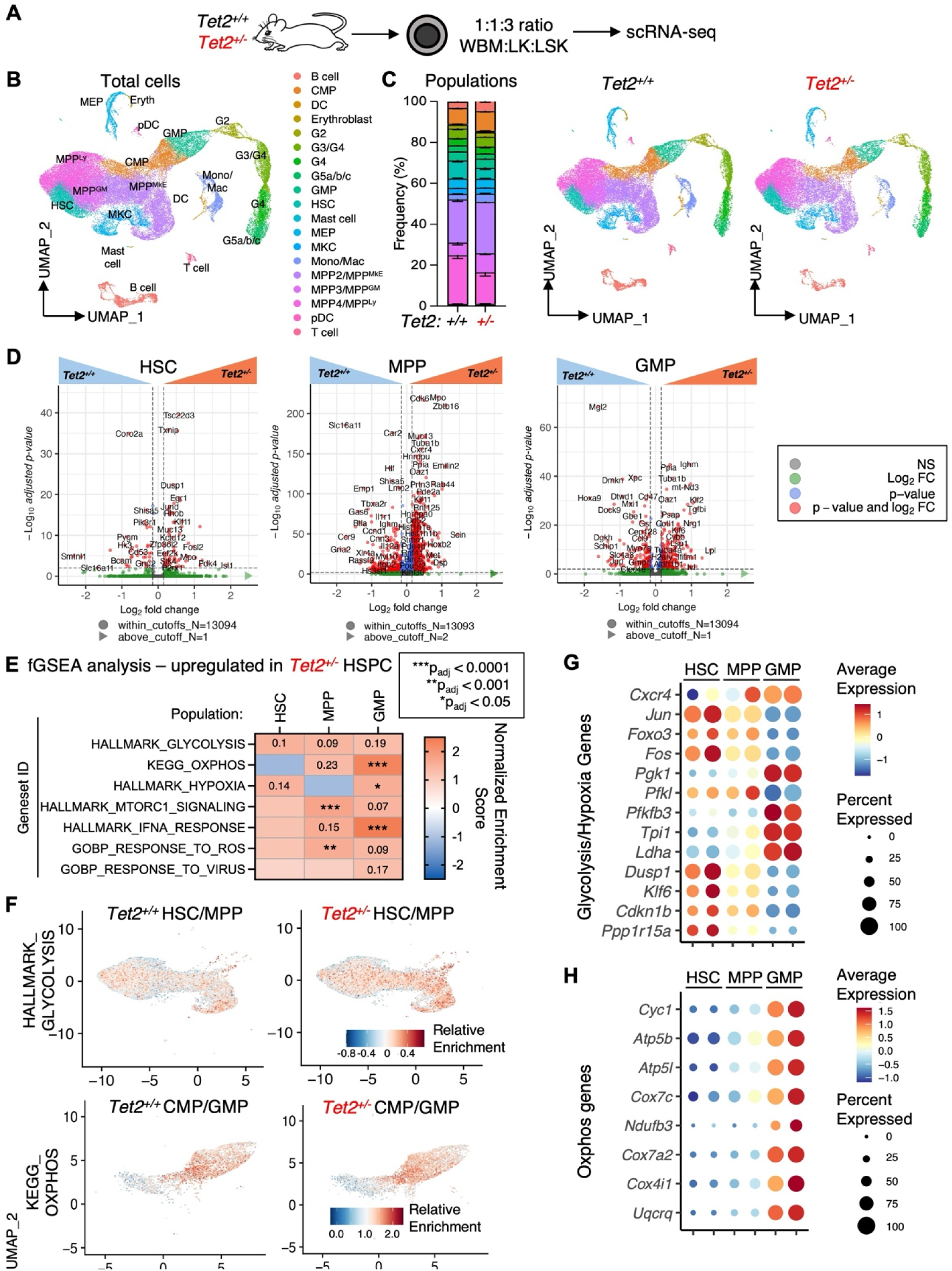
*Tet2*-deficient HSPC overexpress oxidative metabolism gene programs. **A**) Study design showing ratio of BM populations combined for single-cell RNA sequencing (scRNA-seq) analysis; **B**) UMAP projection of scRNA-seq data and resultant population clusters; **C**) relative distribution of populations defined via clustering analysis expressed as frequency of total (left) and UMAP projections separated by genotype; **D**) volcano plots showing differential gene expression between *Tet2^+/+^* and *Tet2^+/-^* HSC (left), combined MPP population (comprised of MPP^MkE^, MPP^GM^, and MPP^Ly^ clusters, center) and GMP (right). Data are expressed as log2 fold change versus -log10 adjusted *p*-value (*p_adj_*) and color-coded to reflect significance. Selected gene symbols are shown on each plot and number of genes is shown below each. Arrow symbols indicate genes above the log2fold change cutoff of the plots. **E**) Heatmap of fast Gene Set Enrichment Analysis (fGSEA) analysis showing pathways upregulated in *Tet2^+/-^*HSC, MPP and GMP. Heatmap scale reflects normalized enrichment score (NES) and asterisks indicate significance based on p_adj_ value. Numbers in heatmap represent p_adj_ values nearing statistical significance. **F**) UMAP projection showing relative enrichment for HALLMARK_GLYCOLYSIS and KEGG_OXPHOS pathways in the indicated combined populations. Dot plots showing percent and average expression of **G**) glycolysis/hypoxia pathway genes and **H**) oxphos pathway genes in HSC, MPP and GMP.

### Increased glucose oxidation is a metabolic feature of *Tet2*-deficient HSPC

To validate and extend our gene expression data, we performed Seahorse flux assays to quantify cellular metabolism in *Tet2^+/+^* and *Tet^+/-^* bead-enriched c-Kit^+^ HSPC (**Fig. 3A**). Strikingly, we observed a significant increase in the maximal respiration rate of *Tet^+/-^* HSPC based on oxygen consumption rate (OCR) measurement. These data indicate *Tet2*-deficient HSPC are capable of higher levels of oxidative phosphorylation (oxphos), the dominant mechanism for ATP generation, than *Tet2^+/+^* HSPC (**Fig. 3B**). Furthermore, *Tet^+/-^* HSPC exhibited a significantly increased extracellular acidification rate (ECAR), indicative of increased glycolytic activity in these cells (**Fig. 3C**). To further characterize the metabolic features of *Tet2^+/-^* HSPC, we performed global metabolomic analysis of *Tet2^+/+^* and *Tet^+/-^* c-Kit^+^ HSPC after 12h of culture using ultra high-performance liquid chromatography coupled to mass spectrometry (UHPLC-MS). We identified 151 unique metabolites (**Table S4**). Sparse partial least squares discriminant analysis (sPLS-DA) and analysis of the top 50 differentially abundant metabolites between *Tet2^+/+^* and *Tet2^+/-^*cells indicated a distinctive metabolic phenotype associated with heterozygous *Tet2* deficiency (**Fig. 3D-E**). Pathway analysis of differentially abundant metabolites identified enrichment for several catabolic pathways associated with oxidative metabolism, including citric acid cycle (also termed tricarboxylic acid, or TCA cycle), transfer of acetyl groups into mitochondria, pyruvate metabolism, Warburg effect (indicative of glucose oxidation), and glutathione metabolism (indicative of redox regulation) (**Fig. 3F**, **Table S5**). Among significantly differentially abundant metabolites detected in our dataset, we observed an increase in metabolites associated with oxidative metabolism in *Tet2^+/-^* HSPC, including ATP, nicotinamide adenine dinucleotide (NAD^+^), which is a coenzyme required for multiple stages of oxidative metabolism, and citrate, which constitutes the entry point of acetyl-CoA into the citric acid cycle (**Fig. 3G**). We independently validated increased ATP abundance using the luminescence-based CellTiter Glo assay (**Fig. 3H**). We also quantified relative NAD^+^/NADH ratios in *Tet2^+/+^*and *Tet2^+/-^* c-Kit-enriched HSPC and identified a relative increase in the NAD^+^/NADH ratio in *Tet2*-deficient cells, consistent with an increase in oxidative metabolism (**Fig. 3H**). We likewise observed a similar pattern of increased ATP levels in sorted HSC/MPP populations after 12h culture (**Fig. 3I**). To assess the impact of gene dosage on metabolic deregulation in the setting of *Tet2* loss of function, we analyzed oxidative flux in *Tet2^+/-^* and *Tet2^-/-^* HSPC versus *Tet2^+/+^*control c-Kit^+^ HSPC using Seahorse analysis (**Fig. S3A**). Expectedly, we found that *Tet2^-/-^* HSPC exhibited proportionally higher oxygen consumption rates than *Tet2^+/-^*cells, indicating loss of both *Tet2* alleles further amplifies metabolic deregulation (**Fig. S3B**). We also analyzed the metabolome of *Tet2^-/-^* c-Kit^+^ HSPC in parallel with our *Tet2^+/-^* and *Tet2^+/+^* cells (**Fig. S3C-D**), and observed similar, though expectedly greater in magnitude, patterns of enrichment for citric acid cycle intermediates (citrate, malate, fumarate) along with NAD^+^ and ATP levels in *Tet2^-/-^* HSPC (**Fig. S3E-F, Tables S4-S5**). These data aligned with published metabolic profiling of *Hoxb8*-immortalized *Tet2*^-/-^ GMPs (35) (**Fig. S3G**). To mechanistically link glycolysis with oxidative metabolism, we performed an independent U-^13^C6 D-glucose stable isotope tracing assay in *Tet2^+/+^*and *Tet2^-/-^* c-Kit^+^ HSPC and measured the fraction of ^13^C isotopologue incorporation into metabolites downstream of glucose (read out as [M+x] with M+0 representing the unlabeled fraction) (**Fig. 3H-I, Table S6**). After 24h of labeling in culture, all intracellular glucose consisted of the ^13^C6 labeled isotopologue, and consistent with increased extracellular acidification rates identified by Seahorse, we observed increased ^13^C labeling of lactate in *Tet2^-/-^* HSPC (**Fig. S3J**). We likewise observed increased ^13^C labeling of TCA cycle intermediates (citrate, malate, fumarate) in *Tet2^-/-^* HSPC (**Fig. S3K**) and into key biosynthetic endpoints, including amino acid (glutamate), fatty acid (acetyl-carnitine), and redox regulation (glutathione) pathways (**Fig. S3L**). Given we observed increases in mitochondrial gene expression in *Tet2^+/-^* HSPC (Fig. 2), to address the biological basis for increased oxidative metabolism, we assessed whether *Tet2*^+/-^ HSPC exhibit changes in mitochondrial membrane potential (Δψ_m_) or mitochondrial abundance. Interestingly, staining of HSPC isolated directly *ex vivo* with tetramethylrhodamine ethyl ester (TMRE) identified no changes in Δψ_m_, in contrast to recently published findings in *Dnmt3a* mutant HSPC (**Fig. 3J**) (36). On the other hand, heterozygous *Tet2*-deficient HSC and MPP exhibited increased staining for the mitochondrial protein Tomm20 (37), indicative of an enlarged mitochondrial network (**Fig. 3K**). This finding is consistent with recently published characterization of mitochondrial DNA levels and electron microscopy analysis of *Hoxb8*-immortalized *Tet2*^-/-^ HSPC (38). Collectively, these findings suggest that loss of *Tet2* triggers increased oxidative metabolism via expansion of the mitochondrial network, versus increased Δψ_m_ associated with *Dnmt3a* loss. To address the importance of increased mitochondrial metabolism in facilitating selective expansion of heterozygous *Tet2-*deficient HSPC, we employed shRNAs targeting essential genes in mitochondrial complex I (*Ndufv1*) and complex IV (*Cox15*) (39). We performed *in vitro* BM competition assays between *Tet2^+/-^* or *Tet2^+/+^* test c-Kit^+^ cells (both CD45.2^+^) and WT CD45.1^+^ c-Kit^+^ competitor cells, combined at a 2:3 test::competitor ratio (**Fig. S3M**). We transduced the mixed BM with shRNAs and following transduction, performed competition assays using serial replating in methylcellulose. Knockdown of both genes substantially reduced the competitive fitness of *Tet2^+/-^* HSPC (**Fig. S3N**). Taken together, these data support a model in which heterozygous *Tet2*-deficient HSPC utilize an expanded mitochondrial network to oxidize glucose at a faster rate than WT HSPC, thereby facilitating their selective advantage.

**Figure 3.**
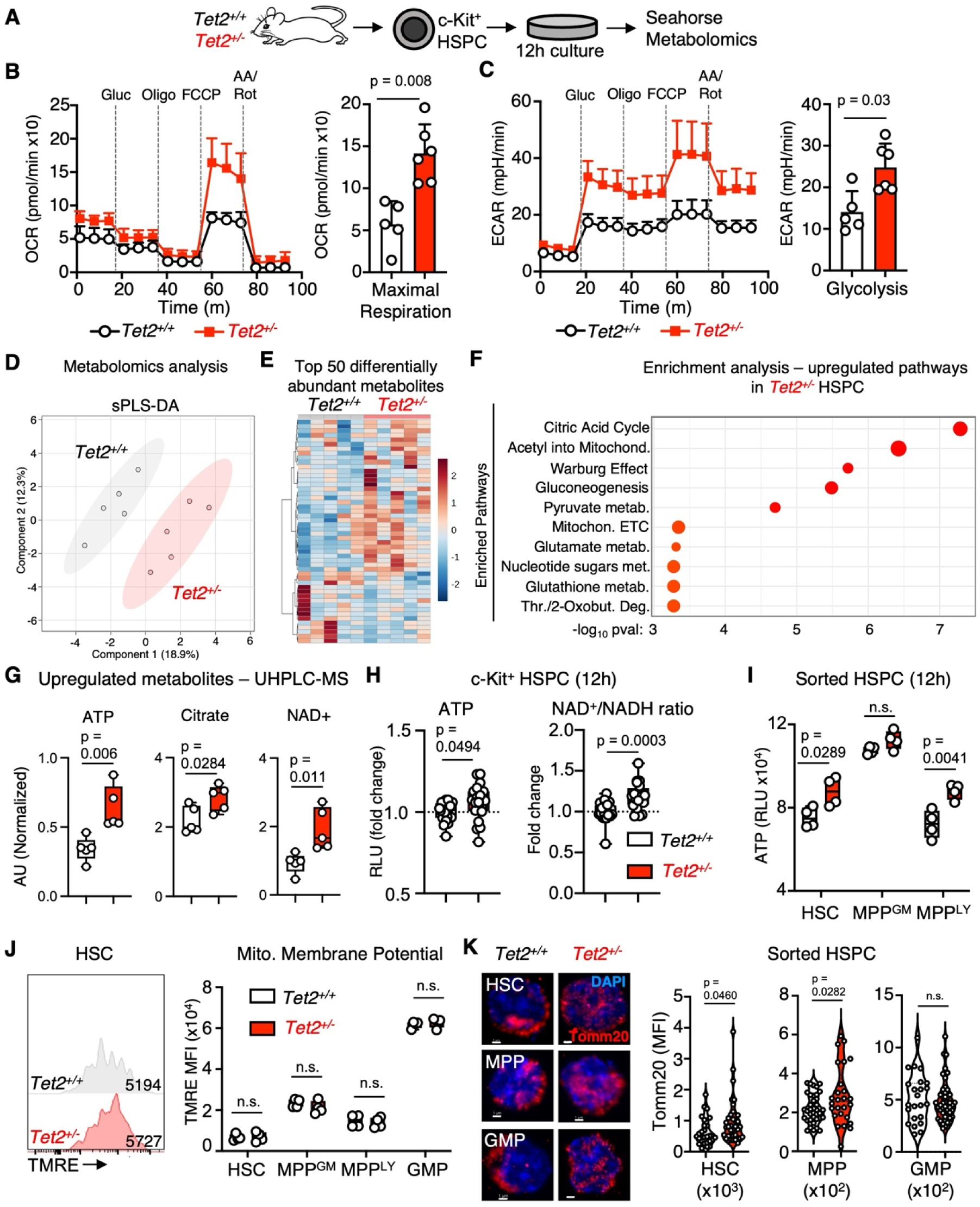
*Tet2*-deficient HSPC exhibit aberrant oxidative metabolism. **A**) Study design. **B**) Mitochondrial stress test of c-Kit+ HSPC cultured for 12h (left) and quantification of maximal respiration (oxygen consumption rate [OCR] after addition of carbonyl cyanide p-trifluoromethoxyphenylhydrazone [FCCP], right); n = 5 *Tet2^+/+^*, 6 *Tet2^+/-^*. Media additions (glucose, olicomycin, FCCP, antimycin A [AA], rotenone) and addition times are shown. **C**) Glycolysis stress test of c-Kit HSPC (left) and quantification of glycolysis (extracellular acidfication rate [ECAR] after the addition of glucose, right); n = 5 *Tet2^+/+^*, 6 *Tet2^+/-^*. **D**) Sparse partial least squares discriminant analysis (sPLS-DA); **E**) hierarchical clustering analysis of top 50 differentially abundant metabolites; **F**) over-representation analysis (ORA) of significantly differentially upregulated metabolites in *Tet2^+/-^* c-Kit^+^ HSPC showing top 10 most significantly enriched metabolic pathways; **G**) quantification of ATP, citrate and NAD^+^ abundance from ultra-high performance liquid chromatography-mass spectrometry (UHPLC-MS) metabolomics analysis of c-Kit+ HSPC cultured for 12h (n = 5/grp). Data in G) are expressed as normalized arbitrary units (AU). **H**) Levels of intracellular ATP and NAD^+^/NADH ratio in c-Kit+ HSPC after 12h culture; **I**) levels of intracellular ATP in sorted HSC and MPP fractions after 12h culture, as read out via luminescence-based assays. **J**) Representative flow cytometry plot (left) and quantification of TMRE fluorescence in the indicated HSPC populations (right); n = 4/grp. Data are expressed as geometric mean fluorescence intensity (MFI) of TMRE. **K**) Representative confocal fluorescence micrographs showing Tomm20 and DAPI staining of GMP (scale bar = 1 μm, left) and quantification of intracellular Tomm20 levels in FACS-sorted HSC (n = 27 *Tet2^+/+^*, 36 *Tet2^+/-^*), MPP (defined as LSK/CD48^+^ HSPC; n = 38 *Tet2^+/+^*, 26 *Tet2^+/-^*) and GMP (n = 23 *Tet2^+/+^*, 47 *Tet2^+/-^*) (right). Data are expressed as mean fluorescence intensity (MFI). Error bars in B) and C) show mean ± SEM. Boxplots show means and individual datapoints and whiskers show minimum/maximum values. Significance was determined via Mann-Whitney u-test or two-way ANOVA with Tukey’s post-test and exact *p*-values are shown. n.s. – not significant. Data in B-C and H-K are compiled from two independent observations. Data in D-G represent one of two independent observations.

### Aberrant oxidative metabolism is a conserved feature of human CH and CCUS

We next sought to validate our results in the setting of human CH. Thus, we leveraged previously published scRNA-seq analyses of BM cells from human subjects with *TET2* mutant CH (**Fig. S4A**). Strikingly, using GSEA we identified significant enrichment for Oxphos and ATP metabolic process gene signatures across hematopoietic ontogeny, including in HSC/MPP fractions, as well as in GMP (**Fig. S4B, Table S7**). To assess whether these features are specific to the mutant HSPC fraction, we likewise reanalyzed published TARGET-seq data (EGAS00001007358) (40), a scRNA-seq modality which allows for discrimination between *TET2* WT and mutant HSPC, using GSEA (**Fig. 4A**). Once again, we found enrichment for Oxphos and ATP metabolic process gene signatures was a specific feature of the *TET2* mutant HSPC fraction (**Fig. 4B, Table S8**). Altogether, these data demonstrate that relative to *TET2* WT HSPC, *TET2* mutant HSPC overexpress oxidative metabolism genes, a feature specific to the mutant HSPC fraction. We next assessed whether these gene expression features are likewise conserved in clonal cytopenia of unknown significance (CCUS), a pathogenic stage of CH that exhibits features of pre-malignant hematopoiesis and carries elevated risk of progression to myelodysplastic syndrome (MDS) (41,42). To examine the gene expression profiles of CCUS HSPC, we performed scRNA-seq analysis of lineage-negative (Lin-) CD34^+^ HSPCs isolated from the BM of three patients with CCUS (**Fig. 4C**, **Table S9**) versus three age-matched healthy donor controls (HC). All patients we profiled carried mutations in *TET2* at a variable variant allelic frequency (VAF), and in some cases also harbored *DNMT3A*, *ASXL1*, *EZH2*, and other CH-associated mutations (**Table S9**). Our scRNA-seq analysis identified 16 cellular clusters (**Fig. 4D**) which we defined based on the differential expression of previously validated lineage-specific transcription factors (TFs) and cellular markers among the clusters (43-46) (**Fig. 4D**, **Fig. S6C**). Differential analysis of cell composition revealed that HSPCs from CCUS patients had a predominantly myeloid-biased differentiation trajectory, which was characterized by an increased frequency of early myeloid/lymphoid progenitors and committed myeloid progenitors, with decreased frequencies of lymphoid progenitors and HSCs (**Fig. 4E-F**). Flow cytometry analysis of samples from a larger cohort of HC and CCUS patient BM specimens showed that CCUS patients have significantly fewer numbers of CD34^+^CD38^-^ HSCs and CD34^+^CD38^+^ hematopoietic progenitor cells in the BM (**Fig. S4D-E**), consistent with clinical presentation of cytopenia in these individuals. Differential analysis of gene expression showed a broad upregulation of metabolic gene programs in CCUS HSPC, including programs related to anabolic processes including cytoplasmic and mitochondrial mRNA translation and proteostasis, as well as neutrophil degranulation and interferon signaling, indicative of immune dysfunction (**Fig. 4G, Tables S10-S11**). Strikingly, we also observed significant upregulation of Oxphos and Glycolysis gene signatures, which underscores the similarity between the metabolic state of HSPCs in human CCUS and CH. Furthermore, we observed across-the-board upregulation of Oxphos genes in CCUS HSPC that were likewise overexpressed in *Tet2*-deficient mouse HSPC fractions (**Fig. 4H**). To validate and extend our gene expression data, we performed ion chromatography-mass spectrometry (IC-MS) analysis of Lin^-^CD34^+^ HSPCs isolated from the BM of CCUS patients and HC (**Table S12**). Using this approach, we identified 132 distinct metabolites. Interestingly, these analyses confirmed that despite genetic heterogeneity among the specimens, CCUS CD34^+^ HSPCs exhibited a distinct metabolic phenotype versus HC, characterized by significant upregulation of intermediates involved in the citric acid cycle pathway (**Fig. 4I-J**, **Fig. S4F, Table S13**). Furthermore, pathway analysis of upregulated metabolites identified enrichment for citric acid cycle, acetyl into mitochondria, and glycolysis (Warburg effect) signatures (**Fig. 4K**).These results suggest that CCUS HSPCs use enhanced oxidative metabolism to meet their energetic needs, likely facilitating their survival and expansion in the BM (47,48). Taken as a whole, these data also demonstrate conservation of dysregulated oxidative metabolism between human CH/CCUS and corresponding mouse models.

**Figure 4.**
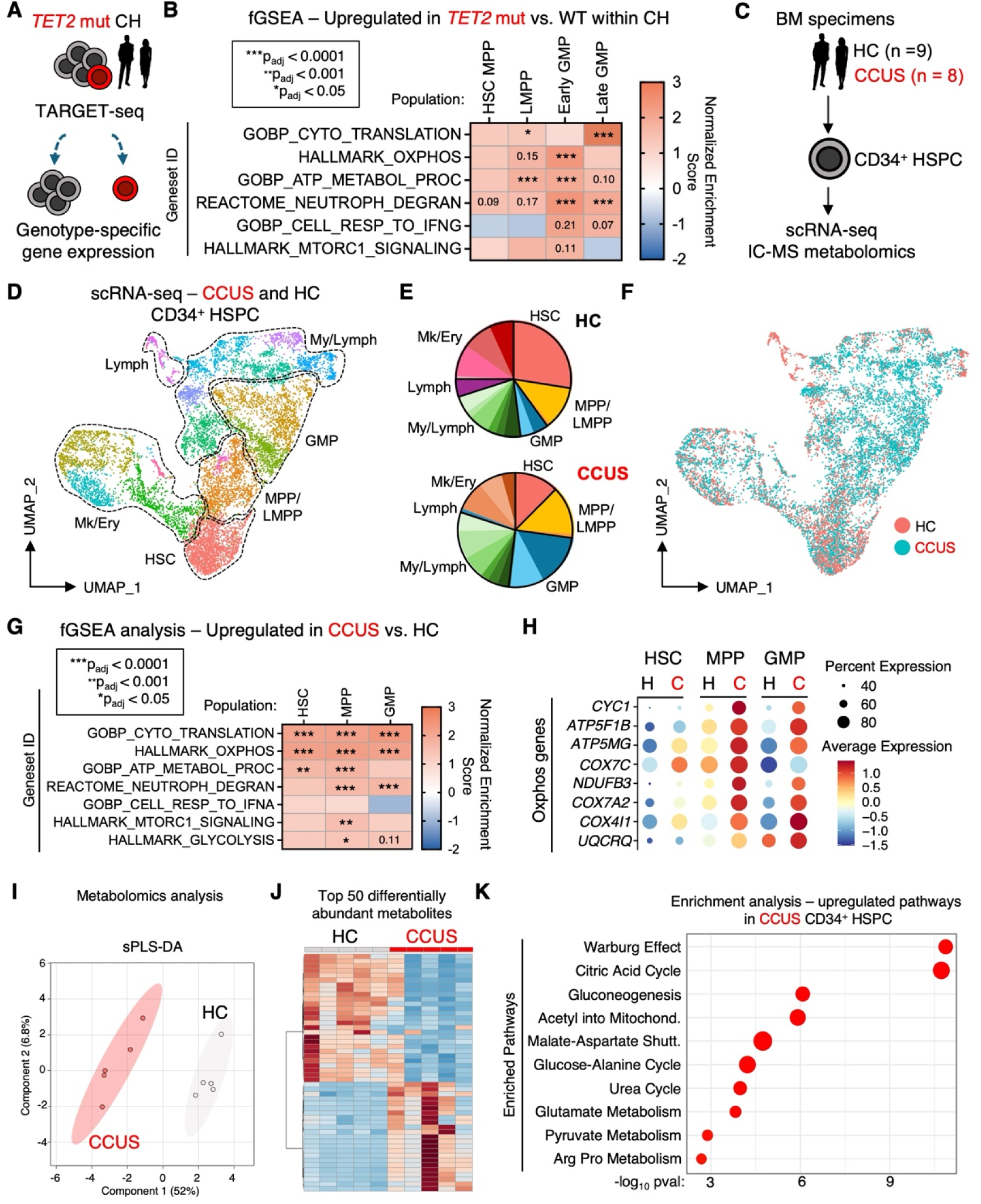
Aberrant oxidative metabolism is a feature of human CH and CCUS. **A**) Study design for published TARGET-seq analysis of human CH subject BM; **B**) Heatmap of Gene Set Enrichment Analysis (GSEA) of combined HSC/MPP, lymphoid MPP (LMPP), early and late GMP clusters. Heatmap scale reflects normalized enrichment score (NES). Asterisks indicate significance relative to *p_adj_* or *p* value. **C**) Study design for scRNA-seq and metabolomics analysis of HSPC from healthy control (HC) and clonal cytopenia of undetermined significance (CCUS) BM specimens; **D**) UMAP projection of scRNA-seq data and resultant population clusters; **E**) relative distribution of populations defined via clustering analysis expressed as frequency of total and **F**) UMAP projection with HC versus CCUS denoted by color. **G**) Heatmap of fast Gene Set Enrichment Analysis (fGSEA) analysis showing pathways upregulated in CCUS HSC, MPP and GMP. Heatmap scale reflects normalized enrichment score (NES) and asterisks indicate significance relative to *p_adj_* or *p* value. **H**) Dot plot showing percent and average expression of Oxphos pathway genes in the indicated populations (H, healthy control; C, CCUS). Ion chromatography-mass spectrometry (IC-MS) analysis of CD34+ HSPC from HC and CCUS BM specimens, n = 5/grp). **I**) Sparse partial least squares discriminant analysis (sPLS-DA); **J**) hierarchical clustering analysis of top 50 differentially abundant metabolites; **K**) over-representation analysis (ORA) of significantly differentially upregulated metabolites in CCUS HSPC showing top 10 most significantly enriched metabolic pathways (n = 5/grp). Data in I-K are compiled from two independent observations.

### *Tet2*-deficient HSPC maintain redox balance despite increased oxidative metabolism

Mitochondrial oxphos triggers production of intracellular reactive oxygen species (ROS), which if in excess can impair HSC self-renewal (49). Given *Tet2*-deficient HSPC exhibit increased oxidative metabolism while retaining enhanced self-renewal capacity, we next quantified intracellular ROS levels in HSPC from our *Tet2*-deficient mouse model. Interestingly, flow cytometry analysis of ROS levels using CellROX dye uncovered no increase in intracellular ROS in FACS-sorted *Tet2*^+/-^ LSK or GMP cells cultured for 12h (**Fig. 5A-C**), suggesting *Tet2*-deficient HSPC can maintain redox homeostasis despite increased levels of oxidative metabolism. ROS produced by mitochondrial respiration is neutralized by an enzymatic cascade in which peroxides are converted to H_2_O by glutathione peroxidase (GPX) via oxidation of glutathione (GSH), resulting in conversion to glutathione disulfide (GSSG), with high glutathione levels being critical for stem cell function (50,51). GSSG is subsequently reduced via glutathione S-reductase (GSR), which in turn oxidizes its cofactor NADPH to NADP^+^. Hence, recycling of glutathione ultimately depends upon the activity of glucose 6-phosphate dehydrogenase (G6PD), the rate-limiting enzyme of the pentose phosphate pathway (PPP), which recycles NADP+ back to NADPH via oxidation of glucose-6P (**Fig. 5D**). Of note, we did not observe broad changes in the expression of glutathione metabolism or PPP pathway genes, including *Gsr* and *Gpx* genes, in our scRNA-seq analyses (**Fig. 5E, Tables S2-S3**), though we did observe modest but significant increases in *G6pdx* expression in *Tet2^+/-^*HSPC (**Fig. S5A**). We therefore interrogated the activity of the glutathione recycling pathway using luminescence-based assays to quantify relative levels of GSH and GSSG in c-Kit^+^ HSPC cultured for 12h (**Fig. 5F**). We observed significantly increased levels of reduced glutathione (GSH) versus glutathione disulfide (GSSG), suggestive of efficient glutathione recycling (**Fig. 5G**). On the other hand, we observed a small but significant shift toward increased NADP^+^ levels versus NADPH, indicative of increased rates of NADPH oxidation needed to facilitate glutathione recycling (**Fig. 5H**). On the other hand, we did not observe changes in the abundance of key metabolites in the non-oxidative PPP, which is not directly involved in control of ROS levels (**Fig. S5B**). Altogether, these data indicate that despite increased rates of oxidative metabolism and a larger mitochondrial network, *Tet2*-deficient HSPC can maintain homeostasis and ROS levels.

**Figure 5.**
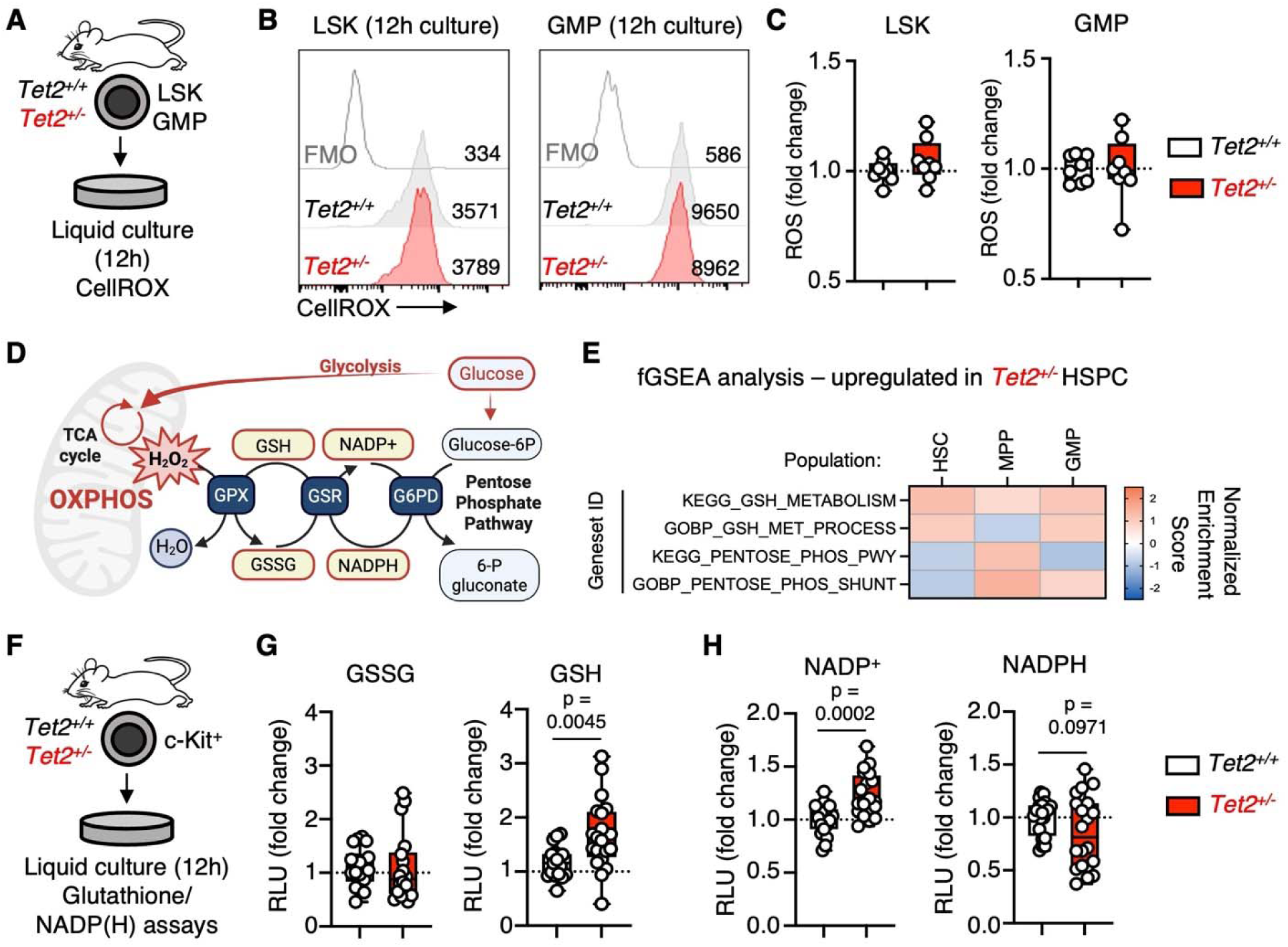
*Tet2*-deficient HSPC maintain redox homeostasis. **A**) Study design. **B**) Representative flow cytometry plots of LSK (left) and GMP (right) CellROX staining after 12h culture (FMO – fluorescence minus one control). Representative geometric mean fluorescence intensities (MFI) are shown in each plot. **C**) Quantification of ROS levels in LSK (left) and GMP (right). Data are expressed as fold change in geometric MFI of CellROX relative to *Tet2^+/+^* cells; n = 8/grp. **D**) Schematic showing ROS detoxification and GSH regeneration pathways. **E**) Heatmap of fast Gene Set Enrichment Analysis (fGSEA) analysis showing pathways upregulated in *Tet2^+/-^* HSC, MPP and GMP. Heatmap scale reflects normalized enrichment score (NES) and asterisks indicate significance relative to *p_adj_* or *p* value. Numbers in heatmap represent *p*-values. **F**) Study design. **G**) Quantification of intracellular GSSG (left) and GSH (right). Data are expressed as fold change in relative luciferase units (RLU) normalized across three independent experiments (n = 20 / grp). **H**) Quantification of intracellular NADP^+^ (left) and NADPH (right). Data are expressed as fold change in relative luciferase units (RLU) normalized across three independent experiments (n = 20 / grp). Boxplots show means and individual datapoints and whiskers show minimum/maximum values. Significance was determined via Mann-Whitney u-test and exact *p*-values are shown. n.s. – not significant. Data in C are compiled from two independent observations. Data in G and H are compiled from three independent observations.

### *Tet2*-deficient HSPC require G6PD to regulate cellular ROS levels

As G6PD is the rate-limiting enzyme of the PPP and directly catalyzes reduction of NADP^+^ into NADPH to facilitate redox homeostasis, we assessed its importance to the function of *Tet2*-deficient HSPC using *in vitro* culture of *Tet2^+/+^* and *Tet2^-/-^*LSK cells with the pharmacological inhibitor 6-aminonicotinamide (6-AN) (**Fig. S5C**) (52). 6-AN suppressed ATP production and selective expansion of *Tet2^-/-^*LSK cells, suggesting G6PD activity is critical for facilitating their metabolic and functional advantage (**Fig. S5D-E**). As 6-AN and other G6PD inhibitors exhibit significant dose-limiting toxicity *in vivo* (53), to address the role of G6PD in maintaining the redox state of *Tet2*-deficient HSPC, we elected to use a genetic approach. Hence, we crossed *Tet2^+/-^* mice with transgenic knockin mice expressing a hypomorphic mutant allele of human *G6PD* (S188F) found in the germline of some individuals of Mediterranean origin (*G6PD^MED^*) (54-56). In humans, this mutation can result in increased sensitivity of red blood cells to oxidative stress and in severe cases, hemolytic anemia. In male mice (which like their human counterparts, are hemizygous due to the location of *G6PD* on the X chromosome) *G6PD^MED^* does not impact steady-state hematological parameters but significantly inhibits erythroid responses to stressors like phenylhydrazine (54). We therefore analyzed the impact of *G6PD^MED^* on hematopoiesis in the BM of male *Tet2^+/+^* and *Tet2^+/-^* mice (**Fig. 6A**). We found no differences in BM cellularity or monocyte abundance across all four genotypes (**Fig. 6B-C**), with only a modest but statistically significant decrease in granulocytes and pre-granulocytes in *Tet2^+/-^*mice. (**Fig. 6C, Fig. S5F**). Interestingly, *Tet2^+/-^* mice harboring the *G6PD^MED^*allele also exhibited a significant increase in GMPs (**Fig. 6D, Fig. S5G**). Despite this, the abundance of HSPC populations was unchanged (**Fig. 6E-G**, **Fig. S5G**). To functionally characterize *G6PD^MED^* HSPC, we analyzed the expansion of LSK in liquid culture for 8d (**Fig. 6H**). Interestingly, *Tet2^+/-^ ::G6PD^MED/-^* LSK cells exhibited a reduced capacity to expand in culture (**Fig. 6I**). This was accompanied by a significant increase in intracellular ROS in *Tet2^+/-^::G6PD^MED/-^* LSK cells versus *Tet2^+/-^* cells after 12h of culture, as determined by CellROX staining (**Fig. 6J-K**). Collectively, these data indicate that *Tet2*-deficient HSPC are highly reliant on the PPP for maintenance of cellular redox state, with G6PD deficiency triggering increased cellular ROS levels and decreased capacity of HSPC to expand in culture.

**Figure 6.**
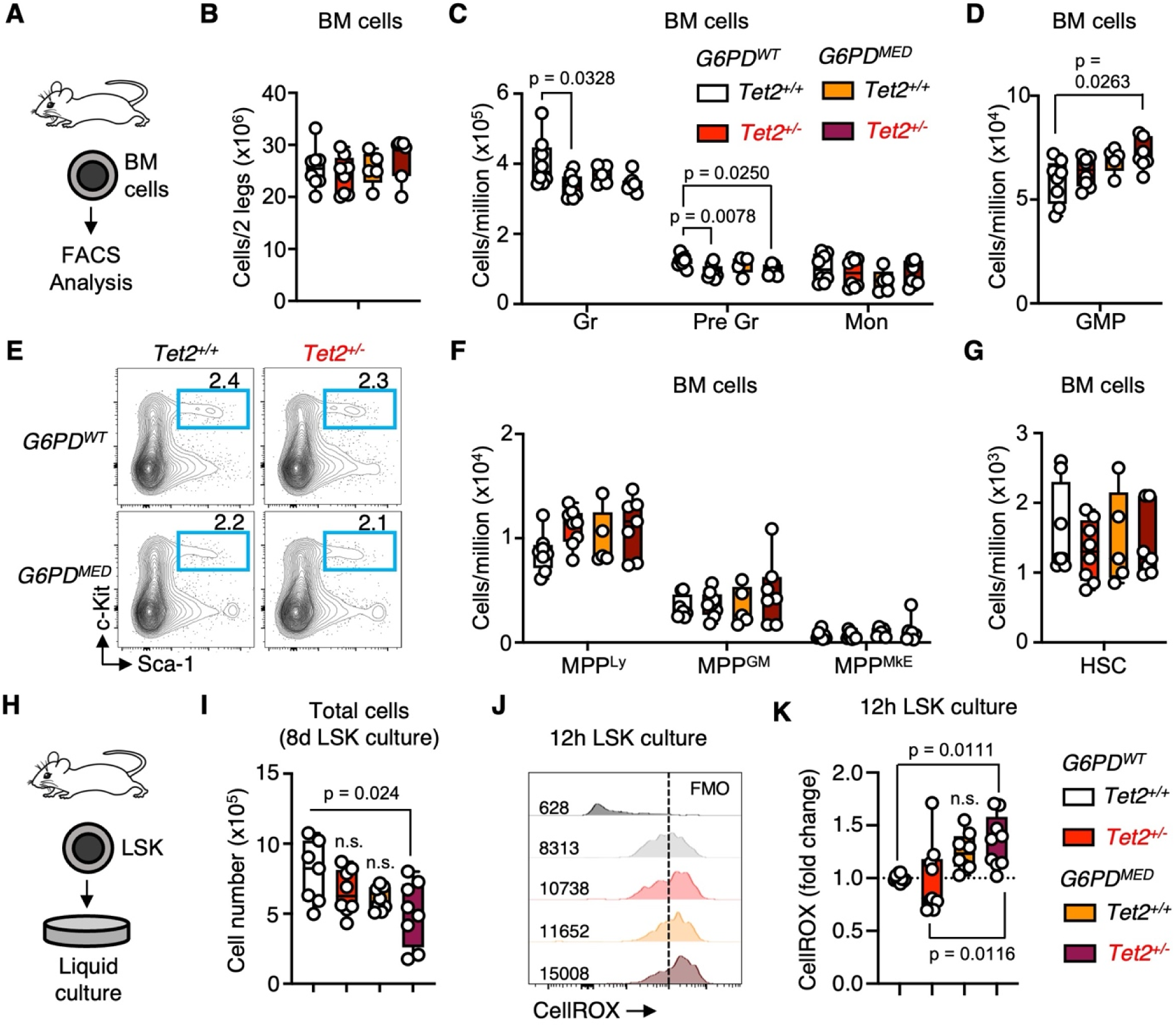
G6PD is required to maintain redox homeostasis in *Tet2*-deficient HSPC. A) Study design (n = 8 *Tet2^+/+^*, *Tet2^+/-^*, n = 5 *G6PD^MED^*; n = 7 *Tet2^+/-^::G6PD^MED^*); **B**) BM cellularity per leg (femur + tibia); quantification of **C**) mature myeloid cells and **D**) GMP in the BM; **E**) representative flow cytometry plots showing gating for LSK populations (HSC/MPP) in BM; **F**) quantification of MPP subtypes and **G**) HSC in the BM. **H**) Study design; **I**) quantification of LSK cell cultures after 8 days (n = 7 *Tet2^+/+^*, *Tet2^+/-^*, *G6PD^MED^*; n = 8 *Tet2^+/-^::G6PD^MED^)*. **J**) Representative flow cytometry plots of LSK cell CellROX staining after 12h culture (FMO – fluorescence minus one control). Representative geometric mean fluorescence intensities (MFI) are shown in each plot. **K**) Quantification of CellROX staining in LSK cells. Data are expressed as fold change in geometric MFI of CellROX relative to *Tet2^+/+^* cells; (n = 10 *Tet2^+/+^*; n = 9 *Tet2^+/-^*and *Tet2^+/-^::G6PD^MED^; n = 8 G6PD^MED^)*. Boxplots show means and individual datapoints and whiskers show minimum/maximum values. Significance was determined via Mann-Whitney u-test or ANOVA with Tukey’s post-test and exact *p*-values are shown. Data in B-G are compiled from three independent observations. Data in I-K are compiled from two independent observations.

### G6PD is required for selective expansion of *Tet2*-deficient HSPC

We next asked whether *Tet2*-deficient HSPC likewise exhibit increased reliance upon G6PD for their selective expansion *in vivo*. Thus, we adoptively transferred BM from all four genotypes of mice (all CD45.2^+^) into non-irradiated CD45.1^+^ recipient mice. We analyzed donor PB chimerism over the course of 20 weeks to assess long-term HSC activity (**Fig. 7A**). Strikingly, we found that *Tet2^+/-^::G6PD^MED/-^*donor chimerism in the PB, while elevated versus *Tet2^+/+^* controls, was significantly reduced vs. *Tet2^+/-^* counterparts (**Fig. 7B-C**). This reduction in donor chimerism was most pronounced in the myeloid lineage, the most direct readout of ongoing HSPC activity (**Fig. 7D-E, Fig. S6A**). Thus, after 24 weeks we analyzed donor chimerism in the BM. Like patterns of donor activity in the peripheral blood, we observed a significant, across-the-board reduction in *Tet2^+/-^::G6PD^MED^*donor chimerism in BM myeloid cell fractions versus *Tet2^+/-^* (**Fig. 7F**, **Fig. S6B**). This phenotype extended into the HSPC compartment, including HSC themselves (**Fig. 7G-I**, **Fig. S6C**). Notably, we did not observe reduced *Tet2^+/+^::G6PD^MED^*donor chimerism in the PB or BM, suggesting that impaired G6PD function on its own is not sufficient to impair HSPC activity. Collectively, our data indicate that loss of G6PD function reduces the capacity of *Tet2*-deficient HSPC to selectively expand in the BM, implicating the PPP as a critical player in regulating aberrant HSPC function in *Tet2* mutant CH. Hence, our data identify loss of PPP function as a metabolic vulnerability that suppresses CH in the setting of *Tet2* deficiency.

**Figure 7.**
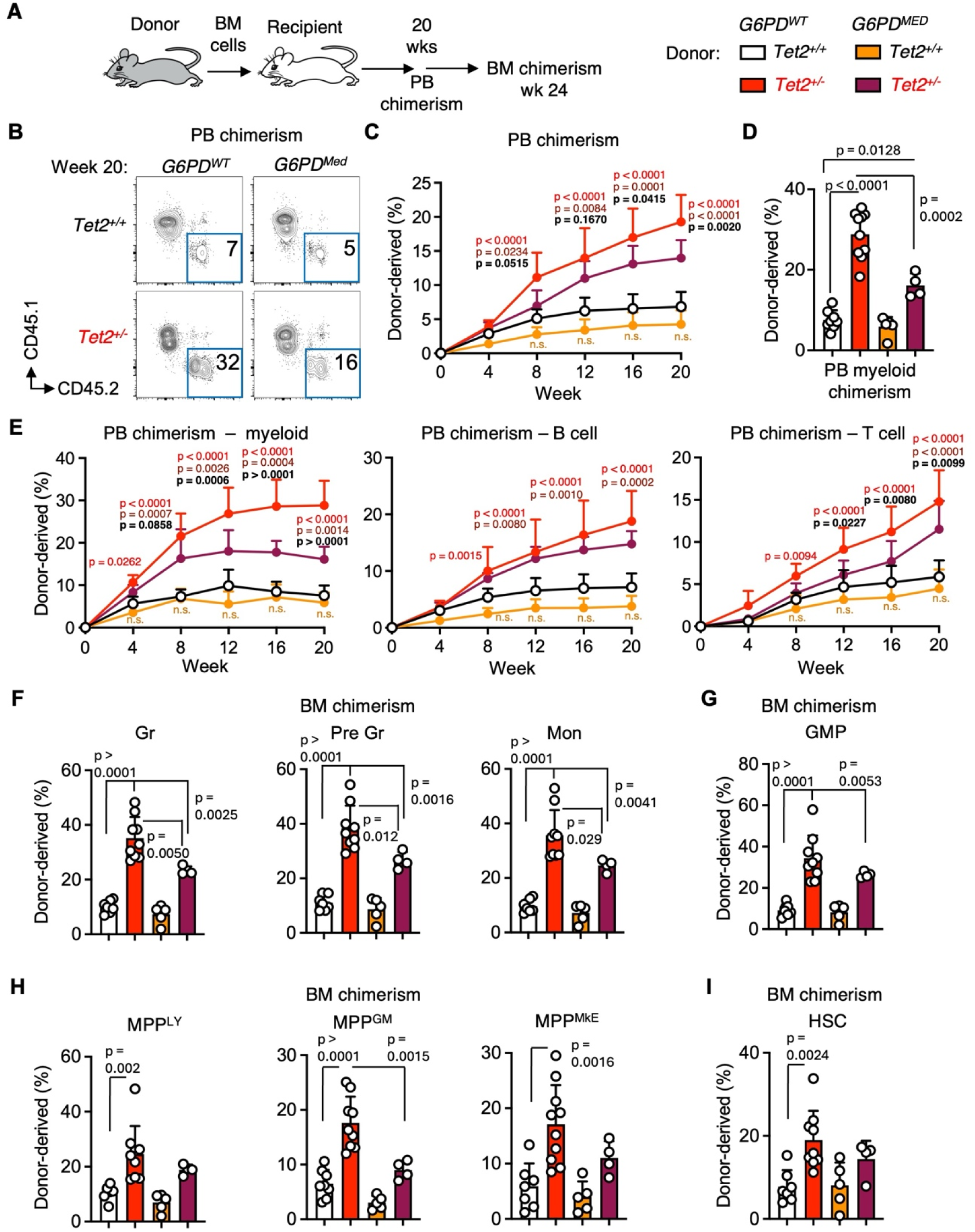
Loss of G6PD activity impairs selective expansion of *Tet2*-deficient HSPC. **A**) Study design (n = 7. *Tet2^+/+^*, n = 9 *Tet2^+/-^*, n = 5 *G6PD^MED^*, n = 4 *Tet2^+/-^::G6PD^MED^*); **B**) representative flow cytometry plots showing donor and recipient PB chimerism at 20 weeks post adoptive transfer; line graphs showing **C**) donor chimerism and **D**) donor chimerism in the CD11b^+^ myeloid compartment at 20 weeks post adoptive transfer; **E**) line graphs showing myeloid (CD11b^+^, left), B cell (B220^+^, center) and T cell (CD3^+^, right) donor chimerism in the PB over time. P-values in red and maroon text represent comparisons between *Tet2^+/+^* and either *Tet2^+/-^* (red) or *Tet2^+/-^::G6PD^MED^* (maroon) genotypes. *P*-values gold represent *G6PD^MED^* genotype. *P*-values in bolded black text represent comparisons between *Tet2^+/-^* and *Tet2^+/-^ ::G6PD^MED^* genotypes. **F**) BM donor chimerism in granulocytes (Gr, left), pre-granulocytes (Pre Gr, center) monocytes (Mon, right); and **G**) granulocyte/macrophage progenitors (GMP); **H**) donor chimerism in lymphoid (Ly), granulocyte/macrophage (GM) and megakaryocyte/erythroid (MkE)-enriched MPP fractions, and **I**) hematopoietic stem cells (HSC) at 24 weeks post adoptive transfer. Error bars represent mean ± SEM. Significance was determined by one-way ANOVA with Tukey’s posttest. Exact *p* values are shown. Data represent one of two independent observations.

## Discussion

CH results from a process of somatic evolution in the BM that facilitates the selective expansion of mutant HSPC clones (1,57). However, the molecular mechanism(s) underlying this process are incompletely understood. Here, we show that *Tet2* deficient HSPC exhibit aberrant overexpression of gene programs that drive oxidative metabolism, licensing them to activate increased levels of oxphos and enhanced production of ATP that facilitates their selective expansion. We show that despite an increased rate of oxidative metabolism, *Tet2*-deficient HSPC maintain nominal levels of ROS, and display a corresponding increase in cellular glutathione (GSH) pools. These data suggest *Tet2*-deficient HSPC are increasingly reliant on G6PD and the PPP to facilitate glutathione reduction and redox homeostasis. We find that HSPC from *Tet2*-deficient mice carrying a hypomorphic allele of *G6PD* exhibit increased levels of ROS and decreased fitness, characterized by significantly decreased donor chimerism versus *Tet2*-deficient controls in a non-conditioned *in vivo* model of CH. Lastly, we show that deregulation of oxidative metabolism is also a feature of *TET2* mutant human CH and CCUS, aligning our mouse studies with human biology. Taken together, our data implicate aberrant cellular metabolism as a driver of selection for *TET2* mutant CH and demonstrate the requirement for cellular redox control via the PPP as a compensatory mechanism to limit the impact of ROS as a byproduct of enhanced oxphos activity.

Over the last decade, the mechanism(s) driving somatic evolution in the BM has become a subject of intensive investigation, as a means of identifying approaches that could be used to limit the potentially harmful consequences of CH. To date, many such studies have focused on characterizing deregulation of transcriptional and epigenetic mechanisms that control HSC fate choices, including the presence of aberrant self-renewal programs that extend the functional capacity of HSC and their progeny in the setting of CH mutations (10,26,58-61). This critical work has identified impairments in key molecular networks controlled by master regulators such as PU.1, CEBPA and other transcription factors (25,35,62,63). While cellular metabolism represents a proximal and highly adaptable regulator of cell fate decisions, relatively few studies have evaluated how CH mutations impact metabolic regulation and in turn, the aberrant functional properties of mutant HSPC.

Here, we find that loss of a single allele of *Tet2* is sufficient to trigger overactivation of glycolysis and mitochondrial oxphos. These findings validate and extend recent work indicating that aberrant mitochondrial metabolism and ATP production in *Tet2*-deficient HSPC is dependent upon ferritinophagy (38). Our investigation identifies cellular redox control via the PPP as a metabolic vulnerability of *Tet2*-deficient HSPC that lies downstream of increased mitochondrial metabolism. In this case, loss of the rate-limiting PPP enzyme G6PD triggers increased ROS and suppresses the selective advantage of *Tet2*-deficient, but not wild-type, HSPC. These data identify aberrant oxidative metabolism and its reactive byproducts as a source of cellular stress in *Tet2*-deficient HSPC that must be buffered, particularly in settings like aging or inflammation where HSPC are apt to become metabolically activated and engage mitochondrial oxphos (64,65). Interestingly, acute myeloid leukemia stem cells (AML LSC) are highly dependent on oxphos for cellular energy production (66,67), and thus inhibition of redox control pathways sensitizes them to cell death via a variety of mechanisms, including impairment of electron transport chain activity and increased oxidative stress (68,69). In contrast to AML LSC, which rely almost exclusively on amino acid metabolism for cellular energy production, we find that *Tet2*-deficient HSPC exhibit elevated levels of glycolysis, which serves as an adaptive mechanism that increases the fitness of *Tet2*-deficient cells via entry into oxidative metabolic pathways. This is potentially relevant as a mechanism of selection in the setting of aging, where normal HSC exhibit reduced glucose uptake (70,71). Further studies can address whether pharmacological targeting of cellular redox potential represents a tractable mechanism to suppress CH clones by limiting their capacity to buffer increased oxidative stress caused by elevated levels of oxidative metabolism.

Aberrant oxidative metabolism may be a convergent phenotype triggered by a variety of molecular and epigenetic deregulations associated with CH mutations and myeloid pathogenesis (72,73). In support of this, three recent studies have identified increased mitochondrial oxidative metabolism as a feature of *Dnmt3a^R882^*-mutant HSPC that contributes to their selective advantage *in vivo* (36,39,74). Numerous factors govern mitochondrial ATP synthesis rates, including the size of the mitochondrial network, Δψ_m_ gradients governing ATP synthase activity, the influx and retention of positively charged ions such as Fe2^+^ and Ca2^+^, mitochondrial quality control mechanisms, and the reductive capacity of antioxidant pathways (37,72,75,76). Of note, in contrast to *Dnmt3a^R882^*mutant HSPC we have not identified increases in Δψ_m_ as the driver of increased oxidative activity in *Tet2*-deficient HSPC. Rather, we observe increases in mitochondrial network size as the key phenotypic feature associated with elevated oxphos activity in *Tet2*-deficient HSPC. Interestingly, the anti-diabetic drug metformin, which among other activities inhibits mitochondrial complex I, has now been identified as a potential therapeutic modality that can suppress selective expansion of *Dnmt3a^R882^* mutant CH(39,74). Retrospective clinical data indicates that the capacity of metformin to suppress CH appears limited to the *Dnmt3a^R882^* mutation (74) and does not impact *TET2* mutant CH, similar to findings in a chimeric *Tet2*-deficient CHIP mouse model treated with metformin (77). Altogether, these results suggest that mechanistically distinct patterns of epigenetic alteration can culminate in deregulated metabolic phenotypes that converge on oxidative metabolism to facilitate increased ATP production and selective advantage. However, they also indicate that the precise mechanism(s) underlying aberrant oxidative metabolism are likely distinct and reflective of the genetic heterogeneity of CH. Targeting redox regulation could thus offer potential for suppressing aberrant oxidative metabolism across multiple genetic mutations by inhibiting a shared compensatory pathway downstream of oxphos that safeguards the aberrant self-renewal of CH clones.

The extent to which other CH mutations, including those in *ASXL1*, splice factors and DNA damage repair pathway factors like *TP53*, culminate in metabolic disruptions that facilitate selection, has not been extensively investigated. Interestingly, our data indicate that relative to HC, CD34^+^ cells from patients with CCUS exhibit features of deregulated oxidative metabolism despite evident genetic heterogeneity among the specimens. Given both *TET2* and *DNMT3A* mutations trigger aberrant oxidative metabolism via distinct molecular and functional pathways, aberrant oxidative metabolism may represent a convergent phenotype that is established early in myeloid pathogenesis and sets the stage for nearly exclusive reliance of malignant stem cells on oxphos at later stages of disease evolution (78). Indeed, recently published work has identified deregulation of the citric acid cycle enzyme COASY as a driver of pathogenesis in *SF3B1* mutant MDS(79), and *TP53* gain of function mutations in LSC likewise can elicit increased oxphos activity (80). We have also found that increased oxphos activity and overexpression of mitochondrial enzymes is a common feature of high-risk MDS HSPC despite heterogenous genetic features (81). Thus, it is likely that aberrant metabolic activity plays a contributory role in selection for multiple CH mutations. Future studies and pre-clinical analyses can identify the spectrum of metabolic phenotypes associated with CH and identify potential strategies to suppress their expansion.

## Methods

### Mice

*Tet2*-deficient mice (B6(Cg)-*Tet2^tm.12Rao^*/J) (Cat# 023359), C57BL/6 mice (Cat#000664) and CD45.1^+^ JaxBoy (C57BL/6J-*Ptprc^em67Lutzy^*/J) mice (Cat# 033076) were obtained from The Jackson Laboratory. G6PD^MED^ mice (54) were a kind gift of Dr. James Zimring (University of Virginia). Mice were housed in a specific pathogen-free animal facility and provided with food and water ad libitum, with a 12-hour light/12-hour dark day schedule. *Tet2*-deficient mice were bred as heterozygotes. *Tet2^+/+^* littermates were used as wild-type controls. 6-18-week-old male and female mice were used for all experiments, except for analysis of G6PD^Med^ and G6PD^Med^::*Tet2^+/-^*mice and respective controls, where only male mice were analyzed due to hemizygosity of male animals for the X-linked *G6PD^MED^* allele. All procedures and treatments were approved by the University of Colorado Anschutz Medical Campus Institutional Animal Care and Use Committee (protocol #00091).

### Patient specimens

BM specimens from patients with CCUS were obtained after the approval of the corresponding Institutional Review Boards (IRB) and in accordance with the Declaration of Helsinki from the Department of Leukemia at MD Anderson Cancer Center (MDACC) under protocol PA15-0926 or from the Department of Medicine at University of California San Diego, BM samples from healthy donors (median age = 54 years, 58% male) were obtained from AllCells (Alameda, CA) or the Department of Stem Cell Transplantation and Cellular Therapy at MDACC. Written informed consent was obtained from all donors, and all diagnoses were confirmed by dedicated hematopathologists. The clinical characteristics of patients with available information are shown in **Table S9**. BM MNCs were isolated from each sample using the standard gradient separation approach with Ficoll-Paque PLUS (GE Healthcare Lifesciences, Pittsburgh, PA). MNCs were enriched in CD34^+^ cells using magnetic separation with the CD34 Microbead Kit (Miltenyi Biotec, San Diego, CA) and further purified or analyzed by flow cytometric sorting.

### Flow cytometry analysis

Flow cytometry analysis was performed in line with previous studies (29,82-84). BM was retrieved from tibiae and femurs of mice via spin flush. One full leg (femur + tibia) was placed in a 0.6 mL microcentrifuge tube that was perforated at the bottom using an 18 G syringe. The 0.6 mL microcentrifuge tube containing femur + tibia was then placed in a 1.7 mL microcentrifuge tube containing 100 uL of staining media (SM, Hanks’ Buffered Saline Solution + 2% Fetal Bovine Serum) and spun at 8000 RPM for 30 seconds to flush out the BM. Cells were then resuspended in 1X ACK solution (150 mM NH4Cl/10 mM KHCO3) for removal of erythrocytes, washed with SM, and filtered through 70-uM mesh to be counted on a ViCell Blu (Beckman Coulter) automated cell counter. All flow cytometry was performed using a Becton Dickenson (BD) FACSCelesta. For all stains described, cells were blocked using 50mg/mL Rat IgG (Sigma). For HSPC analyses, 10^7^ cells were stained in SM on ice for 30 minutes using the following antibodies: PE/Cy5-conjugated anti- CD3, CD4, CD8, Gr1, Ter119, and B220 (as lineage cocktail), Flk2-Biotin, EPCR-PE, CD34-Alexa Fluor 647, CD48-Alexa Fluor 700, and cKit-APC/Cy7. Cells were washed and resuspended in a secondary stain cocktail using 1:4 dilution of Brilliant Staining Buffer (BD) in SM on ice for 30 minutes using the following antibodies: Sca1-BV421, CD41-BV510, Streptavidin-BV605, FcγR-BV711, CD150-BV785. For measurement of donor BM chimerism, CD45.2-FITC and CD45.1-PE/Cy7 were included in the panel. After secondary staining, cells were washed and resuspended in SM containing 1 μg/ml propidium iodide (PI; Sigma-Aldrich) to exclude dead cells. For mature BM and PB population analyses, 10^6^ cells were stained in SM on ice for 30 minutes using the following antibodies: Gr1-PB, Mac1-PE/Cy7, IgM-APC, CD3-Alexa Fluor 700, CD19-APC/Cy7, and Ly6C-Biotin. Cells were washed and resuspended in a secondary stain cocktail using 1:4 dilution of Brilliant Staining Buffer (BD) in SM on ice for 30 minutes using the following antibodies: CD4-BV510, CD8-BV605, Streptavidin-BV711, and B220-BV785. For measurement of donor BM chimerism, CD45.2-FITC and CD45.1 PE were included in the panel. After secondary staining, cells were washed and resuspended in SM containing 1 μg/ml PI to exclude dead cells. A complete list of mouse antibodies is provided in **Table S14**. Quantitative flow cytometry of human live BM MNCs and CD34^+^ cells were performed using a previously described gating strategy and antigen panel (85) that includes antibodies against CD2 (RPA-2.10; 1:20), CD3 (SK7; 1:10), CD14 (MφP9; 1:20), CD19 (SJ25C1; 1:10), CD20 (2H7; 1:10), CD34 (581; 1:20), CD56 (B159; 1:40), CD123 (9F5; 1:20), and CD235a (HIR2; 1:40; all from BD Biosciences); CD4 (S3.5; 1:20), CD11b (ICRF44; 1:20), CD33 (P67.6; 1:20), and CD90 (5E10; 1:10; all from ThermoFisher Scientific, Waltham, MA); CD7 (6B7; 1:20) and CD38 (HIT2; 1:20; both from BioLegend, San Diego, CA); CD10 (SJ5-1B4; 1:20; Leinco Technologies, St Louis, MO); and CD45RA (HI100; 1:10; Tonbo Biosciences, San Diego, CA).

### Mouse BM cell sorting

BM isolation for sorting HSPC populations was performed by crushing femurs, tibiae, hips, humeri, and spines of mice using a mortar and pestle in SM as described previously (29,83,86). Cells were then resuspended in 1X ACK for erythrocyte depletion, washed in SM, and filtered through 70 mm mesh. Cells were separated from debris using Histopaque 1119 gradient centrifugation (Sigma. To enrich for HSPC, cells were labeled with cKit (CD117) beads (Miltenyi) and ran on an AutoMACS Pro (Miltenyi) using the Posseld2 setting. Following enrichment, cells were stained on ice for 30 minutes in SM containing a 1:4 dilution of Brilliant Staining Buffer (Becton Dickinson) using the following antibodies: PE/Cy5-conjugated anti- CD3, CD4, CD8, Gr1, Ter119, and B220 (as our lineage cocktail), Sca1-BV421, FcγR-BV711, CD150-BV785, Flk2-PE, CD48-Alexa Fluor 700, and cKit-APC/Cy7. cells were washed and resuspended in SM containing 1 μg/ml propidium iodide to exclude dead cells. All sorting was performed on a BD Aria Fusion instrument using a 100 mm nozzle. Cells were double sorted to ensure purity.

### Bone marrow adoptive transfer experiments

Adoptive transfer of unfractionated BM into non-irradiated mice was carried out based on established protocols (22). Mouse BM was isolated as described above. and following erythrocyte depletion using ACK, was filtered through a 70 mm filter and resuspended in SM at a concentration of 1 x 10^8^ cells/ml. Mice were anesthetized using isoflurane and 1 x 10^7^ cells (100 ml bolus) were injected retro-orbitally each day for two consecutive days. Donor chimerism was measured in the peripheral blood via flow cytometry as described above.

### Flow cytometry-based assays

For CellRox and MitoSOX assays, LSK and GMPs were isolated from mouse BM as described above. 5 x 10^3^ cells were plated in SILAC media (Gibco) supplemented with the following: dialyzed FBS (5%), HEPES (Gibco, 1mM), Antibiotic-Antimyotic (Sigma, 1X), L-glutamine (Gibco, 1 mM), Arginine (Sigma, 1 mM), Lysine (Sigma, 1 mM), glucose (Sigma, 10 mM), and stem cell factor (SCF; PeproTech, 25 ng/ml). After 12 hrs in culture, cells were transferred over from culture plates to v-bottom 96-well plates, washed with SM, and stained for CellROX/MitoSOX for 30 min at 37°C. For CellROX staining, CellROX Green (Invitrogen) was diluted at 1:10 in DMSO and then added to culture media at 1:500. For MitoSOX staining, MitoSOX Red (Invitrogen) was diluted at 1:1000 in HBSS containing calcium and magnesium. To exclude dead cells from analysis, Live/Dead Fixable Near-IR Dead Cell Stain (Invitrogen) was included in mitochondrial stains at 1:1000. Flow cytometry was performed on a BD FACSCelesta for both CellROX and MitoSOX stains. For TMRE staining, 8 x 10^6^ BM cells were isolated from femurs and tibiae via spin flush and stained on ice in SM containing a 1:4 dilution of Brilliant Staining Buffer (BD) for cell surface markers using 50mg/mL Rat IgG (Sigma) as a block and the following antibodies: PE/Cy5-conjugated anti- CD3, CD4, CD8, Gr1, Ter119, and B220 (as our lineage cocktail), Flk2-Biotin, FcγR-BV711, CD150-BV785, CD34-FITC, Sca1-PE/Cy7, CD49-Alexa Fluor 700, cKit-APC/Cy7. Cells were washed with SM and resuspended in a secondary stain in SM on ice for 30 minutes for Streptavidin-BV605. After the secondary stain, cells were washed in SM and stained with TMRE (Sigma, 10 nM) in 1X DPBS for 30 minutes at 37°C. Cells were then washed in DPBS and resuspended in DAPI (BioLegend, 0.5ug/mL) in DPBS supplemented with 2% FBS for 10 minutes prior to running on a BD Fortessa.

### Confocal microscopy and image analysis

GMP, LSK-CD48+ (MPP) and HSC were sorted and plated for 1 hr in a rectronectin-coated glass slide in the incubator. Cells were then fixed with 4% paraformaldehyde at RT for 10 mins. After washing with PBS, cells were permeabilized with 0.1% Tween 20 for 7 mins at RT. Cells were washed and blocked with phosphate buffered saline (PBS) containing 5% bovine serum albumin (BSA) for 30 mins at RT. After washing, cells were incubated in dark with Tomm20 (ab221292) antibody conjugated to Alexa-555 (1:100) overnight at RT. After washing, the coverslip was mounted in prolong gold antifade mountant containing DAPI (Thermo Fisher Scientific). The slide was cured for 24 hr at RT before imaging. Three-dimensional (3D) confocal imaging was performed using a Nikon A1R inverted LUNV confocal microscope equipped with a 100x oil objective lens. Confocal Z slices of the cells were acquired every 0.2 µm. 3D surfaces were constructed, and mean florescence intensity (MFI) was calculated using Imaris software (Oxford Instruments).

### Luciferase-based metabolic assays

Cells for ATP, NAD+/NADH, NADP+/NADPH, and GSSG/GSH luminescence quantification were isolated as described above. C-Kit+ HSPC were cultured in SILAC media (Gibco) supplemented with the following: dialyzed FBS (5%), HEPES (Gibco, 1mM), Antibiotic-Antimyotic (Sigma, 1X), L-glutamine (Gibco, 1 mM), Arginine (Sigma, 1 mM), Lysine (Sigma, 1 mM), glucose (Sigma, 10 mM), and stem cell factor (SCF; PeproTech, 25 ng/ml). For culture of sorted LSK populations, additional cytokines (thrombopoietin (TPO; 25 ng/ml), IL-3 (10 ng/ml), GM-CSF (20 ng/ml), Flt3L (50 ng/ml), IL-11 (50 ng/ml), and erythropoietin (4 U/ml) from the same supplier were added to the cultures. Cell plating numbers each Glo-assay reaction were as follows: 1 x 10^4^ for ATP, 1 x 10^5^ for total Gluathione/GSSG, 1 x 10^5^ for NAD/NADH, and 1 x 10^5^ for NADP/NADPH. ATP levels were assessed after 12 hours in culture using the CellTiter-Glo assay kit (Promega). For ATP quantification, 100 ml of CellTiter-Glo reagent was added to 100 ml of cell and media suspension in a 96-well opaque plate. 100 ml of media with 100 ml of CellTiter-Glo reagent was used as a background control. The plate was placed on an orbital shaker for 2 minutes and then incubated for 10 minutes at room temperature prior to recording luminescence. Total Glutathione/GSSG levels were assessed after 12 hours in culture using the GSH/GSSG-Glo assay kit (Promega). Cells were gently pipette-mixed 10-15x to disturb the cell monolayer, then split in half, where total glutathione and GSSG was measured. Cells were spun down in 96-well plates for 5 minutes at 1250 rpm. Supernatant was removed by manually pipetting, and cells were resuspended in 25 ml HBSS. 25 ml of total glutathione lysis reagent or oxidized glutathione lysis reagent was added to each resuspended well. The plates were placed on an orbital shaker for 5 minutes. 50 ml of glutathione luciferin generation reagent was added to each well. The plates were placed on an orbital shaker for 2 minutes and then incubated for 30 minutes at room temperature. 100 ml of Luciferin detection reagent was added to each well. 25 ml of HBSS with 25 ml total glutathione or oxidized glutathione, 50 ml of luciferin generation reagent, and 100 ml of luciferin detection reagent was used as a background control. The plates were placed on an orbital shaker for 2 minutes and then incubated for 15 minutes at room temperature prior to recording luminescence. NAD/NADH, and NADP/NADPH levels were assessed after 12 hours in culture using the NAD/NADH-Glo assay kit and NADP/NADPH-Glo assay kit, respectively (Promega). Cells were spun down in 96-well opaque plates for 5 minutes at 1250 rpm. Supernatant was removed by manually pipetting, and cells were resuspended in 50 ml PBS. 50 ml of 0.2N NaOH + 1% DTAB was added to each well. Cells were manually pipette mixed 10-15x and then split in half (50 ml x2) into PCR strip tubes. Samples for NAD^+^ and NADP^+^ readout were acid treated with 25 ml of 0.4N HCL, and then all samples (NAD^+^, NADH, NADP^+^, and NADPH) were heat inactivated in a thermocycler at 60°C for 15 minutes. Samples were allowed to cool to room temperature for 10 minutes after heat inactivation. Acid-treated samples were neutralized with 25 ml of 0.5M Trizma base. Untreated samples (for NADP+ and NADPH readout) were treated with 50 ml of 0.4NHCL/0.5M Trizma base 1:1 mix. 50 ml of treated sample for each readout was transferred into a new 96-well opaque walled plate, and 50 ml of NAD/NADH or NADP/NADPH glo reagent was added to each well. 50 ml of PBS treated like an acid or base treated sample with 50 ml of respective Glo reagent was used as a background control. The plates were placed on an orbital shaker for 2 minutes and then incubated for 180 min at room temperature prior to recording luminescence. All Glo assays were read out using a Synergy H1 microplate reader (Biotek).

### Seahorse metabolic flux assays

cKit+ HSPC were isolated as described above via c-Kit (CD117) microbead labeling of whole BM and enrichment via Posseld2 on a BD AutoMACS Pro instrument (Miltenyi). 2x10^7^ cells were plated for 12 hours in SILAC media (Gibco) supplemented with the following: dialyzed FBS (5%), HEPES (Gibco, 1mM), Antibiotic-Antimyotic (Sigma, 1X), L-glutamine (Gibco, 1 mM), Arginine (Sigma, 1 mM), Lysine (Sigma, 1 mM), glucose (Sigma, 10 mM), and stem cell factor (SCF; PeproTech, 25 ng/ml). After 12 hrs in culture, cells were transferred over from culture plate to Seahorse Cell Culture Microplate (Agilent) treated with Cell-Tak (Corning) at a density of 2x10^5^ cells/well in SeahorseXF DMEM media containing L-glutamine (Gibco, 1mM) and sodium pyruvate (Gibco, 1X). The Seahorse extracellular flux cartridge (Agilent) was calibrated using XF calibrant and prepared according to the manufacturers protocol. For a combined readout of the glycolytic and mitochondrial stress test in the same well, the cartridge was loaded with the following combination of reagents: Glucose (Sigma, 100 nM, 20 uL, port A), Oligomycin (Sigma, 20 uM, 22 uL, port B), FCCP (Sigma, 20 uM, 25 uL, port C), and Rotenone/Antimycin-A (Sigma, 5uM, 27 uL, port D). For Mitochondrial Stress Test analysis of *Tet2^+/+^*, *Tet2^+/-^* and *Tet2^-/-^*HSPC, 5x10^4^ cells/well were used to ensure sufficient dynamic range to read out elevated metabolic flux in all three genotypes. Cells were cultured in IMDM (Wisent) with 10% FBS, SCF (100 ng/ml), TPO (25 ng/ml) and Flt3L (50 ng/ml) at 37C for 2hr. Cells were washed 1x and equilibrated for 1hr at 37°C in Seahorse XF DMEM. Only Oligomycin, FCCP and Rotenone/Antimycin-A were included in the cartridge. All Seahorse runs were performed on a Seahorse XFe96 Analyzer, and all metabolic parameters were quantified using calculation software provided by Agilent.

### shRNA preparation and transduction

Viral particles were produced and transduced as previously described (38). HEK293T cells cultured in DMEM +10% FBS and 2mM Gluta Plus were seeded at a density of 7x106 cells in a 15 cm plate 24h prior to transfection. Cells were co-transfected using jetPRIME (Polyplus) according to the manufacturer’s instructions with the psPAX2 packaging plasmid, the VSV-G envelope plasmid, and a pRSI9 plasmid encoding shScramble, shNdufv1, or shCox15 targeting sequences. Supernatants from co-transfected HEK293T cells was collected 48 and 72 hours later, filtered through a 0.45 mm PVDF filter to remove debris, and precipitated overnight in 40% w/v polyethylene glycol (PEG). Viral particles were collected by centrifugation at 3700 RPM for 30 minutes in a refrigerated centrifuge at 4°C, and the viral pellet was resuspended in HBSS + 25 mM HEPES and stored at -80°C.

### Mouse HSPC *in vitro* cell competition assays

Assays were performed as previously described (39). CD45.2^+^ *Tet2^+/+^*or *Tet2^+/-^* c-Kit^+^ test cells were mixed with CD45.1^+^ c-Kit^+^ cells at a 2:3 ratio and the mixture was transduced with lentivirus containing *Ndufv1*, *Cox15* or a scrambled control shRNA (described above). Cells were plated on retronectin-coated plates (Takara, T100B) for 2h at room temperature, blocked with PBS + 2% BSA for 30 minutes at room temperature, and spin infected with virus suspension at 3700 RPM for 2h at 4°C. The combined mixture was subsequently cultured in MethoCult M3434 medium (Stemcell Technologies) at a density of 200 cells per well in a 96-well plate for 7 days. Colonies from replicate wells were harvested, pooled, recounted and serially replated in Methocult at a density of 500 cells per well for another 7 days. Cells were harvested, washed with 1x PBS, stained for CD45.1 and CD45.2 along with SYTOX Green (Thermo Fisher) to exclude dead cells, and analyzed on a CytoFLEX (Beckman Coulter) flow cytometer.

### scRNA-seq library preparation

For mouse specimens, a 3:1:1 ratio of FACS-sorted LSK, FACS-sorted c-Kit^+^ progenitors, and unfractionated BM cells were combined to a total of 3 x 10^5^ cells per sample using a hemocytometer to verify sorter-based cell counts. Samples were spun down at 400 x g for 5 min in a 4°C centrifuge. Following removal of supernatant via pipet, cells were fixed in 1ml of 4% formaldehyde Fixation Buffer from the Chromium Next GEM Single Cell Fixed RNA Sample Preparation Kit following manufacturer’s protocols (10X Genomics). Subsequently, whole transcriptome probe pairs were added to the fixed single cell suspensions and incubated overnight, followed by washing, addition of Quenching Buffer and storage at - 80°C. Single cell suspensions were then loaded onto a Chromium X (10X Genomics) microfluidics instrument for capture. Resulting droplet suspensions were placed in a thermal cycler for 1hr at 25°C, 45 min at 60°C, and 20 min at 80°C. Barcoded products were subjected to library preparation for sequencing as paired-end 150 bp reads on the Illumina NovaSeq 6000 (Illumina, San Diego, CA) at a depth of 20,000 reads per cell.

For human specimens, FACS-purified live Lin^-^CD34^+^ cells were sequenced at the Advanced Technology Genomics Core facility at MDACC. Sample concentration and cell suspension viability were evaluated using a Countess II FL Automated Cell Counter (ThermoFisher Scientific) and manual counting. Samples were normalized for input onto the Chromium Single Cell A Chip Kit (10x Genomics, Pleasanton, CA), in which single cells were lysed and barcoded for reverse-transcription. The pooled single-stranded, barcoded cDNA was amplified and fragmented for library preparation. Pooled libraries were sequenced on a NovaSeq6000 SP 100-cycle flow cell (Illumina, San Diego, CA).

### scRNA-seq analysis

Fastq files were processed using 10x Genomics Cell Ranger v7.1.0 multi pipeline and aligned to the 10x Genomics Chromium Mouse Transcriptome Probe Set v1.0.1. Cell Ranger-filtered data in .h5 format were imported and processed with a Seurat v4.1.1 framework (87). Doublets were classified with scDblFinder v1.8.0 (88) in each sample’s filtered feature barcode matrix prior to merging into a single Seurat object. Genes with counts in less than 10 cells were removed and the log2 genes, log2 UMIs, and the percent of mitochondrial probe counts were quantified by sample and per cell. Low quality cells were flagged within each sample with Scater v1.4.0 (89). A threshold of three median absolute deviations (MADs) above or below the median of the log2 UMI or log2 genes per cell or above the median percent mitochondrial were defined as outliers and subsequently removed. To remove doublets, the remaining cells were subjected to unsupervised clustering analysis and the proportion of doublets per cluster was calculated. Only cells classified as singlets within clusters with lower than three MADs of the median doublet proportion per cluster were kept. A total of 59,182 cells and 13,095 features remained for downstream analysis. The gene counts were divided by each cell’s total counts and multiplied by 10,000 followed by natural-log transformation. The top 2,000 most variable genes were selected with the variance stabilizing transformation method. B- and T-cell receptor features were removed from the set according to published methods (90,91). The remaining 1,986 highly variable genes were scaled and input to PCA. The top 30 principal components were input to UMAP reduction and shared nearest-neighbors clustering analysis using a resolution of 0.9. Selection of the clustering resolution was guided by clustree analysis (92) in conjunction with viewing canonical cell type gene expression profiles measured at neighboring resolutions.

Clusters were identified as cell types through multiple methods. The top 200 enriched genes by cluster were used as input to over-representation analysis (93) with gene sets from the MSigDB C8 collection (94), the PanglaoDB (95), and published cell type signatures of HSC/MPP (32,96) and Granulocyte (97) subtypes. Gene expression of canonical cell type markers was plotted in UMAP space and as dotplots. HSC/MPP and Granulocytes were subset and subtyped separately to account for minor expression differences.

For human BM specimens, sequencing analysis was carried out using 10X Genomics’ CellRanger software, version 3.0.2. Fastq files were generated using the CellRanger MkFastq pipeline (version 3.0.2). Raw reads were mapped to the human reference genome (refdata-cellranger-GRCh38-3.0.0) using the CellRanger Count pipeline. Multiple samples were aggregated using the Cellranger Aggr pipeline. The digital expression matrix was analyzed with the R package Seurat (version 5.2.1) (98) to identify different cell types and signature genes for each. Cells with fewer than 500 unique molecular identifiers or 100 genes or greater than 25% mitochondrial expression were excluded from further analysis. The Seurat function NormalizeData was used to normalize the raw counts. Variable genes were identified using the FindVariableFeatures function. The ScaleData function was used to scale and center expression values in the dataset, and the number of unique molecular identifiers was regressed against each gene. Uniform manifold approximation and projection (UMAP) was used to reduce the dimensions of the data, and the first two dimensions were used in the plots. The FindClusters function was used to cluster the cells. Marker genes for each cluster were identified using the FindAllMarkers function. Cell types were annotated based on the marker genes and their match to canonical markers (43-46).

### Differential Gene Expression analysis

For mouse samples, gene expression differences between groups were measured within cell populations with at least 50 cells in each group using the Wilcoxon rank sum test and multiple testing controlled with the Bonferroni procedure implemented in the FindMarkers function. Significant differentially expressed genes (DEGs) were identified as having an adjusted p-value less than 0.01 and an average log2 fold change greater than 0.15 magnitude in either direction. Two types of pathway enrichment analysis were performed in parallel with select gene sets from the MSigDB GOBP (99,100), KEGG (101,102), Hallmark (103) collections, and custom gene signatures. First, significant DEGs were rank ordered by average log2 fold change and the top 200 DEGs by cluster in either direction were input to separate over-representation (93) analyses. Second, all genes input to differential expression testing were rank ordered by average log2 fold change and subjected to fGSEA analysis. Signature scores were calculated per cell using the Seurat AddModuleScore function and distributions were viewed in UMAP space and by violin plot. In human samples, nominal p-values were corrected by Benjamini-Hochberg procedure and DEGs were identified using an adjusted *p*-value threshold of 0.05 or lower. Pathway enrichment analysis was performed using the Reactome, GOBP, KEGG and Hallmark collections.

### Ultra-High Performance Liquid Chromatography-Mass Spectrometry (UHPLC-MS)

Mouse c-Kit+ cells were isolated by magnetic enrichment as indicated above and cultured. For metabolic labeling studies, cells were first cultured in SILAC Advanced DMEM/F-12 medium (ThermoFisher) with 5% dialyzed FBS, 10mM HEPES, 1mM arginine, 200nM lysine and 10mM U-^13^C6-glucose (Sigma-Aldrich) containing 25ng/ml SCF for 24h. For global (non-labeled) metabolomics analysis, c-Kit+ cells were cultured in the same media formulation as above, with unlabeled D-glucose substituted for ^13^C-labeled glucose, for 12h. Following culture, cells were washed with ice-cold PBS and pelleted by centrifugation at 500 × *g* for 5 min at 4°C. All liquid supernatant was carefully removed via pipetting, and cell pellets were snap frozen using liquid nitrogen. Subsequently, cells were extracted into an ice-cold methanol:acetonitrile:water buffer (at a 5:3:2 v/v/v ratio) to a final concentration of 2 x 10^6^ cells/ml. Specimens were vortexted for 30 min at 4°C and then centrifuged at 12,000 x g for 10 min at 4°C. Supernatants containing extracted metabolites were then subjected to UHPLC-MS. 10 microliters of extract was loaded onto a Kinetex XB-C18 column (150 x 2.1 mm i.d., 1.7±mm – Phenomenex). Metabolites were eluted using a 5 min gradient (Phase A – water + 0.1% formic acid; Phase B – acetonitrile + 0.1% formic acid for positive ion mode; for negative ion mode, Phase A – 5% acetonitrile + 5 mM ammonium acetate; Phase B – 95% acetonitrile + 5 mM ammonium acetate). The mass spectrometer was set to scan in Full MS mode at 70,000 resolution, 65-975 m/z range, 4 kV spray voltage, 45 sheath gas and 15 auxiliary gas. Positive and negative ion analyses were conducted using separate runs. Metabolite assignment was performed using an in-house standard library as a reference. Data analysis and pathway enrichment analysis of significantly differentially abundant metabolites was performed using Metaboanalyst 6.0 (metaboanalyst.ca) (104,105) using default parameters.

### Ion Chromatography-Mass Spectrometry (IC-MS)

Human CD34^+^ cells were isolated by magnetic enrichment as indicated above, washed with ice-cold PBS, and pelleted by centrifugation at 1,000 × *g* for 2 min at 4°C. Cell pellets were snap-frozen in liquid nitrogen and temporarily stored at -80°C. Sample analysis was carried out in the Metabolomics Facility at MDACC. Metabolites were extracted from cell pellets by adding 1 mL of ice-cold 0.1% ammonium hydroxide in 80/20 (v/v) methanol/water. Extracts were centrifuged at 17,000 × *g* for 5 min at 4°C, and supernatants were transferred to clean tubes and subjected to evaporation to dryness under nitrogen. Dried extracts were reconstituted in deionized water, and 10 μL was injected for analysis by ion chromatography–mass spectrometry. The ion chromatography mobile phase A (weak) was water, and the mobile phase B (MPB; strong) was water containing 100 mM KOH. Ion chromatography was performed with the Dionex ICS-5000^+^ system (ThermoFisher Scientific), which included a Thermo IonPac AS11 column (4-µm particle size, 250 × 2 mm) with the column compartment kept at 30°C. The autosampler tray was chilled to 4°C. The mobile phase flow rate was 360 µL/min, and the gradient elution program was 0-5 min, 1% MPB; 5-25 min, 1-35% MPB; 25-39 min, 35-99% MPB; 39-49 min, 99% MPB; 49-50 min,

99-1% MPB. The total run time was 50 min. For better sensitivity, desolvation was assisted by methanol, which was delivered by an external pump and combined with the eluent via a low dead volume mixing tee. Data were acquired using a Thermo Orbitrap Fusion Tribrid Mass Spectrometer under ESI negative ionization mode at a resolution of 240,000. Raw data files were imported into Thermo Trace Finder software for final analysis. The relative abundance of each metabolite was normalized by live cell number. Levels of 1,3-BPG and 3-PG were inferred from the amount of their isomers 2-phospho-glycerate and 2,3-diphosphoglycerate. Data analysis and pathway enrichment analysis of significantly differentially abundant metabolites was performed using Metaboanalyst 6.0 (metaboanalyst.ca) using default parameters.

### Statistical analysis

Statistical analysis was performed using GraphPad Prism 10 software (GraphPad, La Jolla, CA). Statistical tests used are identified in the figure legends. *P*-values are shown in each figure. *P*-values less than or equal to 0.05 were considered statistically significant.

### Data Availability

The data generated in this study are available within the article and its supplementary data files. Expression profile data are publicly available in Gene Expression Omnibus (GEO) at GSE314014 (murine) and GSE310705 (human). Metabolomics datasets are publicly available via Metabolomics Workbench as study numbers ST004480 (murine global metabolomics), ST004481 (murine ^13^C glucose tracing) and ST006532 (human targeted metabolomics). Published TARGET-seq raw data used for independent validation are available at the European Genome-phenome Archive (EGA) at accession EGAS00001007358. Published bulk RNA-seq analysis of murine *Tet2^+/+^*, *Tet2^-/-^*and *Tet2^Mut^* LSK cells (27) used for independent validation is available in GEO at GSE132090.

## Acknowledgements

This project was supported by the Cleo Meador and George Ryland Scott Endowed Chair in Hematology, CU Department of Medicine Outstanding Early Career Scholars Program, R01 DK 137183, American Cancer Society Research Scholar Award RSG-22-138-01-IBCD, and Blood Cancer United (formerly Leukemia and Lymphoma Society) Scholar Award 1392-24 (to E.M.P.), a Blood Cancer United Career Development Program Award 3456-26 (to S.B.P.), an HHMI Gilliam Fellowship and the KUH ASCEND U2C/TL-1 Training Program (to K.E.N.), and by philanthropic contributions to MD Anderson’s AML and MDS Moon Shot Program (to S.C.). This project was supported by a grant to the University of Colorado Cancer Center (P30 CA046934) and by the Alpine HPC system, which is jointly funded by the University of Colorado Boulder, the University of Colorado Anschutz, Colorado State University, and the National Science Foundation (award 2201538). This work used MD Anderson’s South Campus Flow Cytometry and Cell Sorting Core Laboratory and Sequencing and Microarray Facility, supported in part by the NCI through MD Anderson’s Cancer Center Support Grant (P30 CA16672).

## Author Contributions

Conceptualization: V.A., I.G.G., S.C., E.M.P.; Methodology: K.E.N., V.A., A.E.G., M.H., S.B.P., S.M., S.N.B., D.S., M.D.F., G.H., S.X., S.M.C., L.T., I.G.G., A.D.; Investigation: K.E.N., V.A., C.M.C., W.E.S., S.N.B., M.H., S.B.P, S.M., H.Y.L, T.Y., D.S., K.S.C., I.G.G.; Formal Analysis: A.E.G., E.P.D., F.M., C.C.A., D.S., A.V., J.A.R., L.T., A.D.; Data Curation: A.E.G., E.P.D., C.C.A., A.V., K.I., M.M.D., J.A.R., S.X., L.T.; Resources: S.M.C., K.I., M.M.D., S.X., G.G.M., K.S.C., A.D.; Writing – Original Draft Preparation: K.E.N., V.A., A.E.G., C.M.C., S.N.B., S.C., E.M.P.; Supervision: J.A.R., M.D.F, A.D, S.X., S.M.C., G.G.M., S.C., E.M.P.; Funding Acquisition: S.C., E.M.P.

