## Supplementary Figures and Captions for "Aberrant oxidative metabolism selects for *TET2*-deficient hematopoietic stem and progenitor cells"

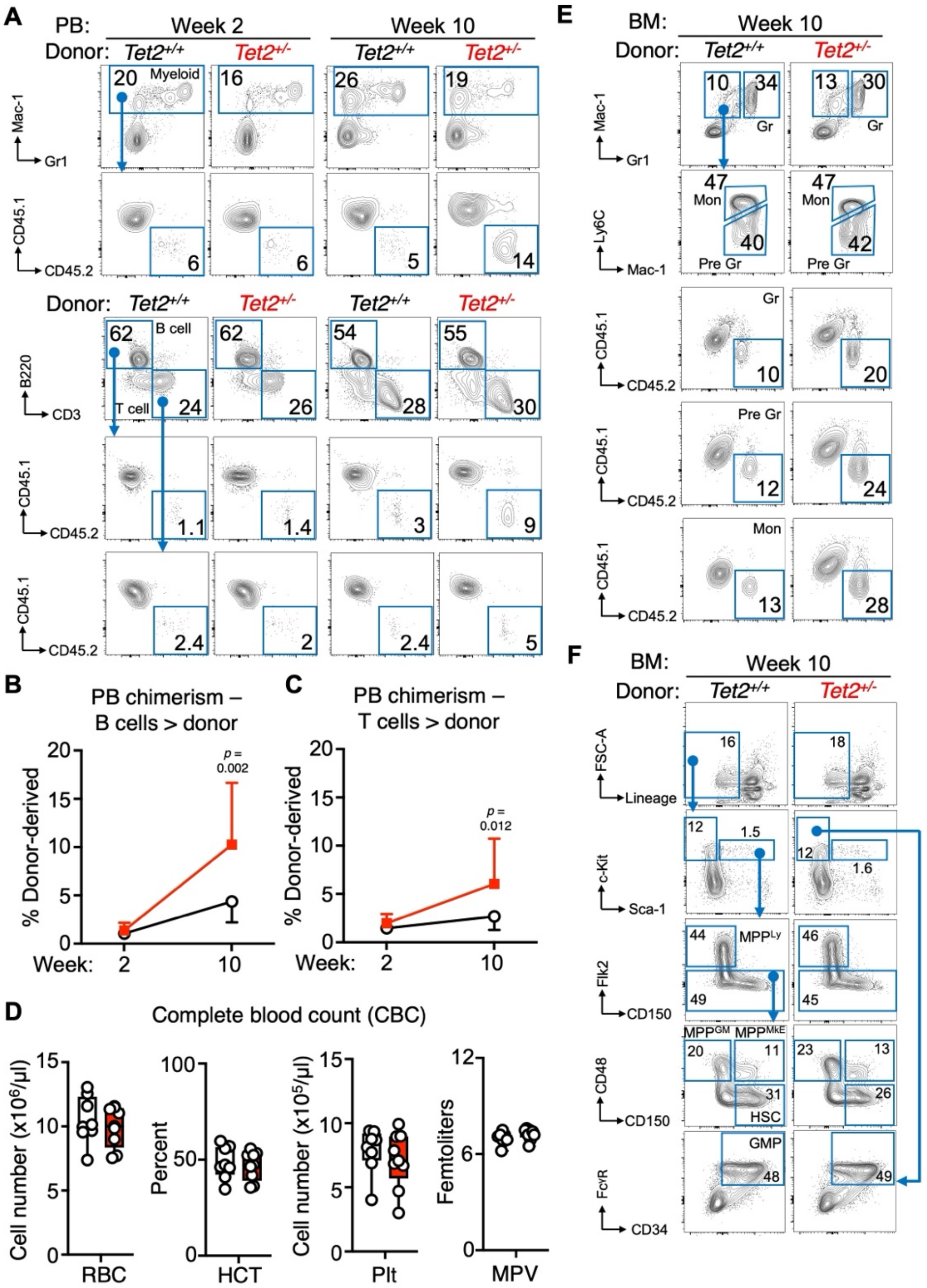


**Figure S1. Flow cytometry analysis of donor chimerism**, related to Figure 1. **A**) Representative flow cytometry plots and gating strategy for analysis of donor chimerism in peripheral blood (PB) of mice in Fig. 1A-D. Representative frequencies within each gate are shown. Line graphs showing **B**) B cell and **C**) T cell donor chimerism in the PB at the indicated times. **D**) CBC analysis of mice in Fig. 1E of red blood cells (RBC), hematocrit (HCT), platelets (Plt), and mean platelet volume (MPV). Representative flow cytometry plots and gating strategy for analysis of donor chimerism in **E**) BM myeloid populations and **F**) BM HSPC populations of mice in Fig. 1F-J. Representative frequencies within each gate are shown. Error bars in B and C represent mean ± SEM. Boxplots show means and individual datapoints and whiskers show minimum/maximum values. Significance was determined via Mann-Whitney u-test or ANOVA with Tukey's post-test and exact *p*-values are shown.

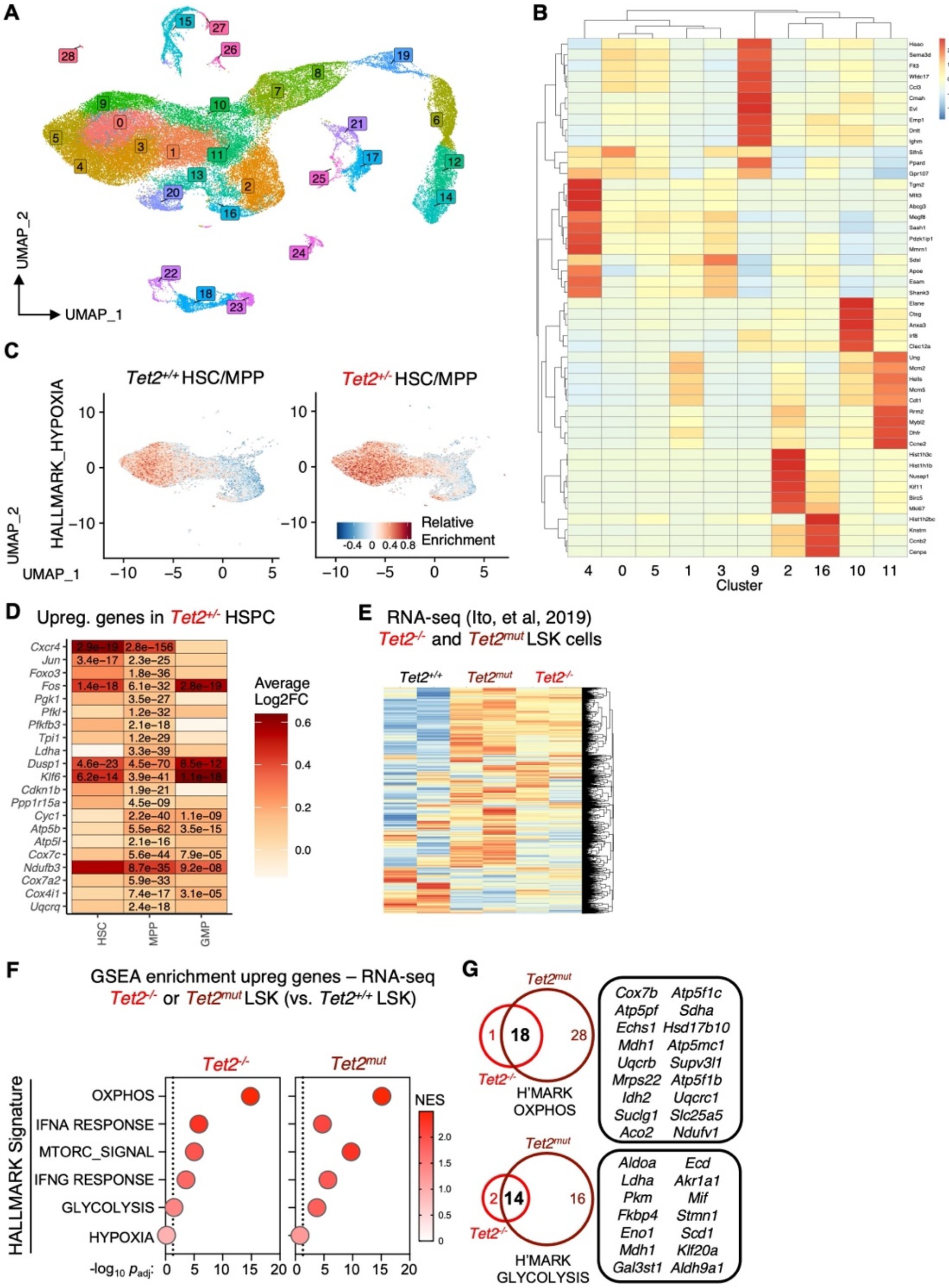


**Figure S2. Clustering of scRNA-seq data and independent validation in bulk RNA-seq**, related to Figure 2. **A**) UMAP projection showing clustering analysis and **B**) heatmap showing expression of genes used for identification of HSPC clusters in scRNA-seq analysis shown in Fig. 2. **C**) UMAP projection showing relative enrichment for HALLMARK_HYPOXIA pathway in the indicated combined populations. **D**) Heatmap showing Log2 fold change for upregulated genes shown in Fig. 2G-H. $p_{adj}$ values are shown for genes where $p_{adj} < 0.05$. **E**) Heatmap analysis of *Tet2*$^{+/+}$, *Tet2*$^{mut}$ and *Tet2*$^{-/-}$ LSK cell bulk RNA-seq data from GSE132090 used as independent validation of scRNA-seq dataset; **F**) Gene Set Enrichment Analysis (GSEA) showing enriched upregulated pathways in *Tet2*$^{-/-}$ (left) and *Tet2*$^{mut}$ (right) LSK cells versus *Tet2*$^{+/+}$ controls. Data are expressed as -log10 $p_{adj}$ with normalized enrichment score (NES) depicted by color intensity. **G**) InteractiVenn analysis of overlapping upregulated genes from the indicated HALLMARK gene signatures between *Tet2*$^{-/-}$ and *Tet2*$^{mut}$ LSK cells.

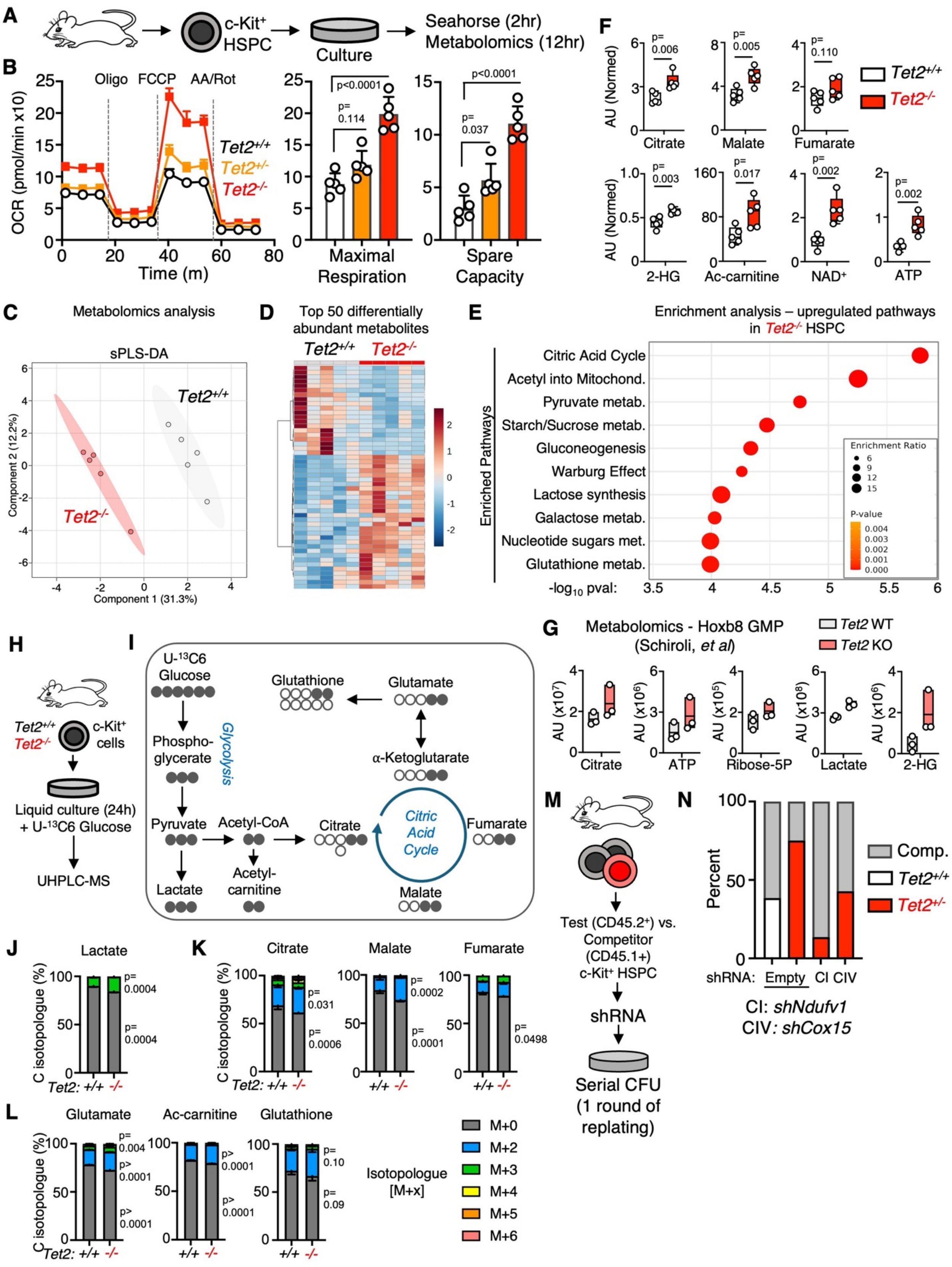
A
c-Kit+ HSPC
Culture
Seahorse (2hr)
Metabolomics (12hr)
B
Oligo FCCP AA/Rot
OCR (pmol/min x10)
Time (m)
Tet2+/+
Tet2+/-
Tet2-/-
p<0.0001
p= 0.114
Maximal Respiration
p<0.0001
p= 0.037
Spare Capacity
F
AU (Normed)
p= 0.006
Citrate
p= 0.005
Malate
p= 0.110
Fumarate
Tet2+/+
Tet2-/-
p= 0.003
2-HG
p= 0.017
Ac-carnitine
p= 0.002
NAD+
p= 0.002
ATP
C
Metabolomics analysis
sPLS-DA
Tet2+/+
Tet2-/-
Component 2 (12.2%)
Component 1 (31.3%)
D
Top 50 differentially abundant metabolites
Tet2+/+
Tet2-/-
E
Enrichment analysis – upregulated pathways in Tet2-/- HSPC
Enriched Pathways
Citric Acid Cycle
Acetyl into Mitochond.
Pyruvate metab.
Starch/Sucrose metab.
Gluconeogenesis
Warburg Effect
Lactose synthesis
Galactose metab.
Nucleotide sugars met.
Glutathione metab.
Enrichment Ratio
P-value
-log10 pval:
G
Metabolomics - Hoxb8 GMP (Schiroli, et al)
Tet2 WT
Tet2 KO
AU (x10^7)
Citrate
AU (x10^6)
ATP
AU (x10^5)
Ribose-5P
AU (x10^8)
Lactate
AU (x10^6)
2-HG
H
Tet2+/+
Tet2-/-
c-Kit+ cells
Liquid culture (24h) + U-13C6 Glucose
UHPLC-MS
I
U-13C6 Glucose
Glutathione
Glutamate
Phospho-glycerate
Glycolysis
α-Ketoglutarate
Pyruvate
Acetyl-CoA
Citrate
Citric Acid Cycle
Fumarate
Lactate
Acetyl-carnitine
Malate
M
Test (CD45.2+) vs. Competitor (CD45.1+) c-Kit+ HSPC
shRNA
Serial CFU (1 round of replating)
N
Percent
Comp.
Tet2+/+
Tet2+/-
shRNA: Empty CI CIV
CI: shNdufv1
CIV: shCox15
J
Lactate
C isotopologue (%)
p= 0.0004
p= 0.0004
Tet2: +/+ -/-
K
Citrate
p= 0.031
p= 0.0006
Malate
p= 0.0002
p= 0.0001
Fumarate
p= 0.0498
L
Glutamate
p= 0.004
p> 0.0001
p> 0.0001
Ac-carnitine
p> 0.0001
p> 0.0001
Glutathione
p= 0.10
p= 0.09
Isotopologue [M+x]
M+0
M+2
M+3
M+4
M+5
M+6

**Figure S3. Metabolomics analysis of *Tet2$^{-/-}$* HSPC**, related to Figure 3. **A**) Study design; **B**) Mitochondrial stress test analysis comparing respiratory activity of cultured *Tet2$^{+/+}$*, *Tet2$^{+/-}$* and *Tet2$^{-/-}$* c-Kit$^{+}$ HSPC (left) and quantification of maximal respiration (oxygen consumption rate [OCR] after addition of FCCP, center) and spare respiratory capacity (maximal respiration – basal respiration, right); n = 5/grp. Media additions and addition times are shown. **C**) Sparse partial least squares discriminant analysis (sPLS-DA); **D**) hierarchical clustering analysis of top 50 differentially abundant metabolites; **E**) over-representation analysis (ORA) of significantly differentially upregulated metabolites in *Tet2$^{-/-}$* c-Kit$^{+}$ HSPC showing top 10 most significantly enriched metabolic pathways; **F**) quantification of significantly upregulated metabolites from ultra-high performance liquid chromatography-mass spectrometry (UHPLC-MS) metabolomics analysis of c-Kit$^{+}$ HSPC cultured for 12h; n = 5/grp. Data in F) are expressed as normalized arbitrary units (AU). Boxplots show means and individual datapoints. Whiskers depict minimum and maximum values. UHPLC-MS analysis of *Tet2$^{-/-}$* HSPC was performed in parallel with analyses of *Tet2$^{+/+}$* and *Tet2$^{+/-}$* HSPC shown in Fig. 3A. **G**) Re-analysis of published metabolomics data from Hoxb8-immortalized GMP. Data are expressed as arbitrary units (AU). Boxplots show means and individual datapoints. **H**) Study design, n = 4 *Tet2$^{+/+}$*, 3 *Tet2$^{-/-}$*; **I**) schematic representation of U-$^{13}$C6 glucose labeling through glycolysis and one turn of the citric acid cycle, along with entry into fatty acid and glutamate/glutathione synthesis pathways. $^{13}$C is depicted as closed circles, $^{12}$C (unlabeled) is depicted as open circles. Quantification of $^{13}$C isotopologue labeling as a proportion of total carbon in **J**) lactate, **K**) TCA cycle and **L**) downstream glucose-derived biosynthetic pathways. Data are shown as stacked bars depicting means $\pm$ SEM for each C isotopologue. **M**) Study design for *in vitro* cell competition assays following shRNA knockdown of *Ndufv1* and *Cox15*, n = 1/grp. **N**) Quantification of proportion of test::competitor cells following one round of serial replating. Significance was determined by ANOVA with Tukey's post test. Exact p values are shown.

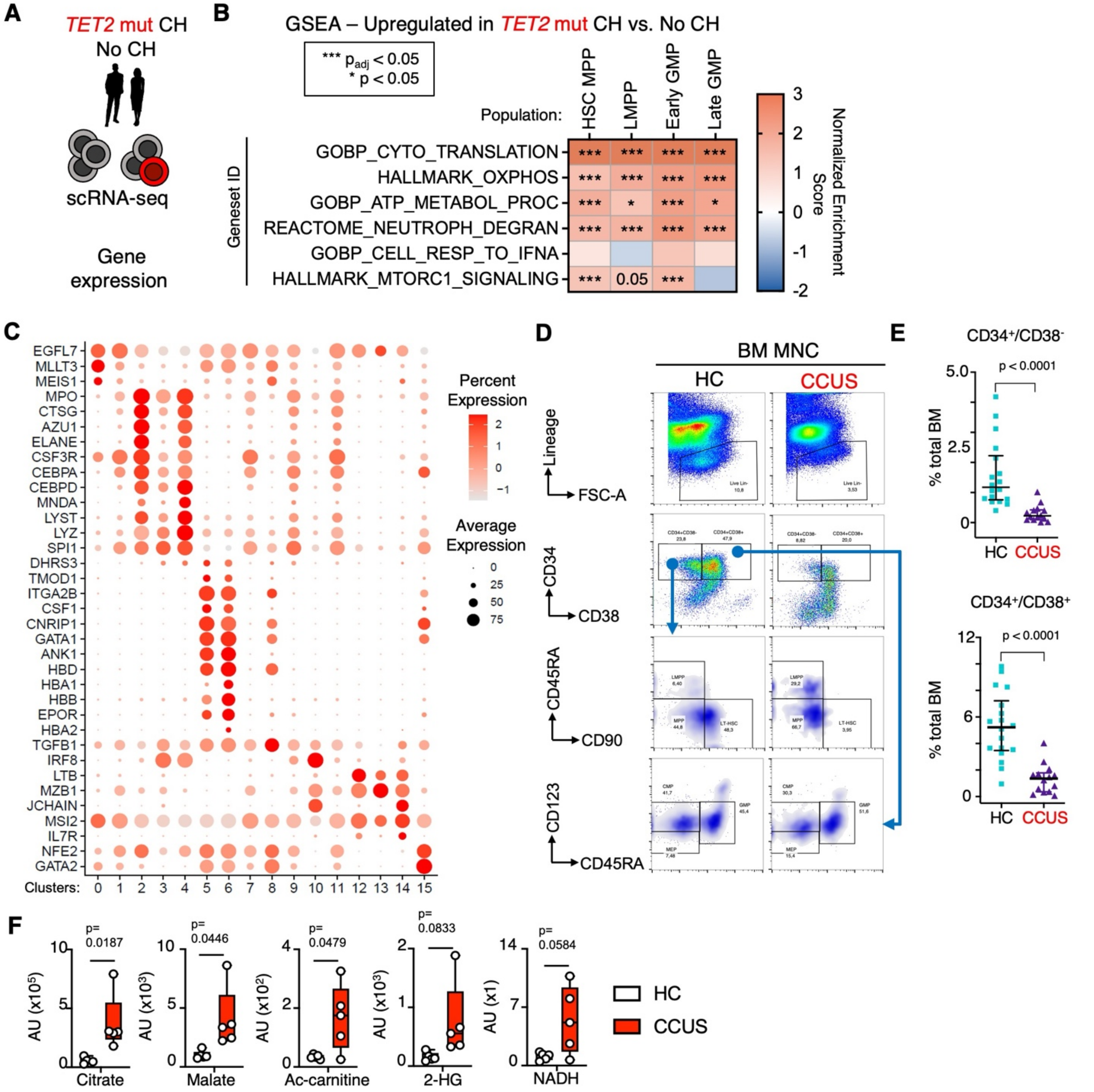


**Figure S4. Clustering of scRNA-seq data and flow cytometry profiling of CCUS HSPC**, related to Figure 4**. A**) Study design for published scRNA-seq analysis of human CH subject BM; **B**) Heatmap of Gene Set Enrichment Analysis (GSEA) of combined HSC/MPP, lymphoid MPP (LMPP), early and late GMP clusters. Heatmap scale reflects normalized enrichment score (NES). Asterisks indicate significance relative to $p_{adj}$ or $p$ value. **C**) Dotplot showing expression of genes used for identification of HSPC clusters in scRNA-seq analysis shown in Fig. 7D. **D**) Representative flow cytometry gating and **E**) quantification of HSPC populations in healthy control (HC) and clonal cytopenia of undetermined significance (CCUS) BM specimens, n = 19 HC, n = 16 CCUS. Individual datapoints are shown, error bars represent mean $\pm$ SEM. **F**) quantification of significantly upregulated metabolites from ion chromatography-mass spectrometry (IC-MS) metabolomics analysis of HC and CCUS HSPC; n = 5/grp. Data in F) are expressed as normalized arbitrary units (AU). Boxplots show means and individual datapoints. Whiskers depict minimum and maximum values. Exact p values are shown.

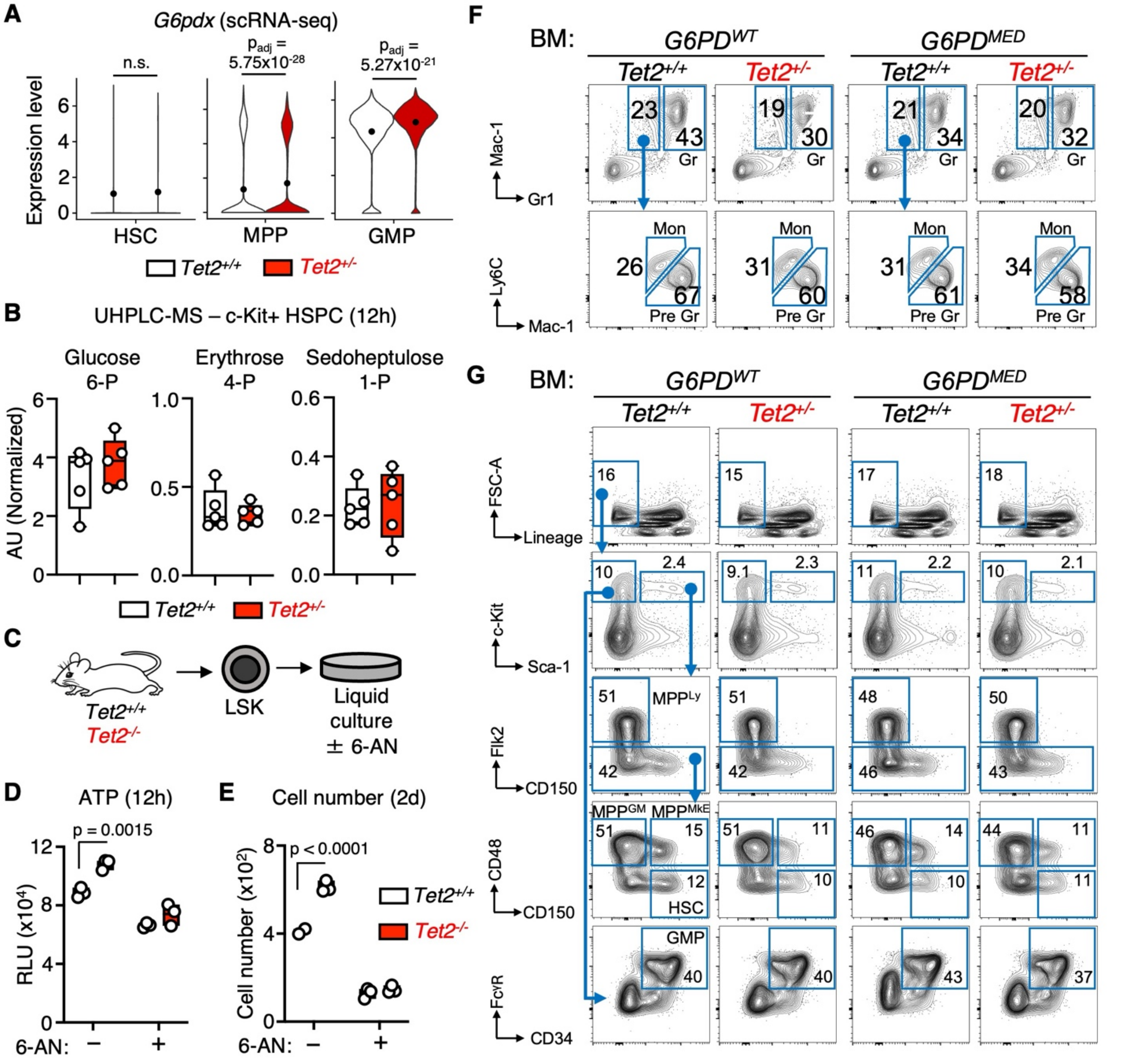


**Figure S5. NADPH levels, G6PD expression and flow cytometry analysis of *G6PD^MED* mice**, related to Figures 5 and 6. **A**) Analysis of scRNA-seq data from Figure 2 for *G6pdx* expression. **B**) Quantification of non-oxidative pentose phosphate pathway (PPP) metabolites by UHPLC-MS in c-Kit$^+$ HSPC from Fig. 3. Data are expressed as normalized arbitrary units (AU). Boxplots show means and individual datapoints. Whiskers depict minimum and maximum values. n = 5 / grp. **C**) Study design for *in vitro* 6-aminonicotinamide (6-AN) G6pd blockade experiment; **D**) quantification of ATP in LSK cells after 12h culture. Data are expressed as relative luciferase units (RLU), n = 3 / grp. **E**) Quantification of cell numbers after 2d culture, n = 3/grp. Boxplots show means and individual datapoints. Representative flow cytometry gating for **F**) mature myeloid and **G**) HSPC populations shown in Figure 6C-G. Representative population frequencies are shown within each gate. Significance was determined by Mann-Whitney u-test or ANOVA with Tukey's posttest. Exact *p* values are shown.

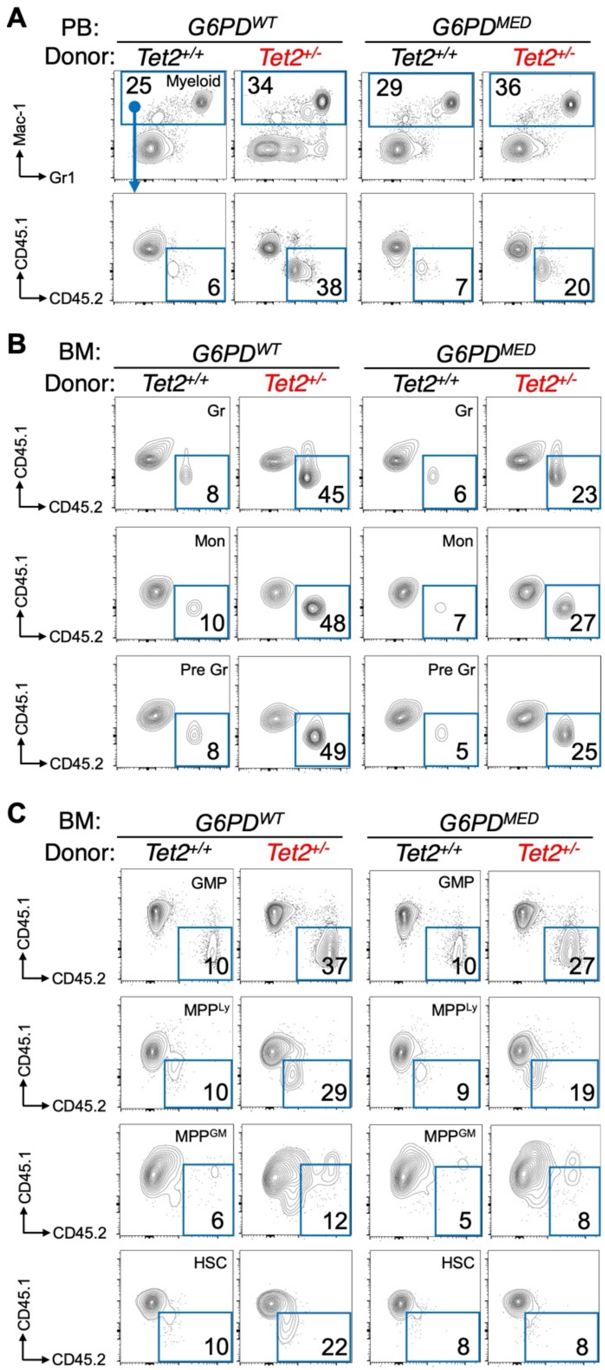


**Figure S6. Flow cytometry analysis of *G6PD$^{MED}$* donor chimerism**, related to Figure 7. **A**) Representative flow cytometry plots and gating strategy for analysis of myeloid donor chimerism in peripheral blood (PB) of mice in Fig. 6A-E. Representative frequencies within each gate are shown. Representative flow cytometry plots and gating strategy for analysis of donor chimerism in **B**) BM myeloid populations and **C**) BM HSPC populations of mice in Fig. 6F-I Representative frequencies within each gate are shown. Gating strategies for identification of BM myeloid and HSPC populations are identical to those shown in Fig. S1E-F.
